# Imputation of ancient genomes

**DOI:** 10.1101/2022.07.19.500636

**Authors:** Bárbara Sousa da Mota, Simone Rubinacci, Diana Ivette Cruz Dávalos, Carlos Eduardo G. Amorim, Martin Sikora, Niels N. Johannsen, Marzena Szmyt, Piotr Włodarczak, Anita Szczepanek, Marcin M. Przybyła, Hannes Schroeder, Morten E. Allentoft, Eske Willerslev, Anna-Sapfo Malaspinas, Olivier Delaneau

## Abstract

Due to *postmortem* DNA degradation, most ancient genomes sequenced to date have low depth of coverage, preventing the true underlying genotypes from being recovered. Genotype imputation has been put forward to improve genotyping accuracy for low-coverage genomes. However, it is unknown to what extent imputation of ancient genomes produces accurate genotypes and whether imputation introduces bias to downstream analyses. To address these questions, we downsampled 43 ancient genomes, 42 of which are high-coverage (above 10x) and three constitute a trio (mother, father and son), from different times and continents to simulate data with coverage in the range of 0.1x-2.0x and imputed these using state-of-the-art methods and reference panels. We assessed imputation accuracy across ancestries and depths of coverage. We found that ancient and modern DNA imputation accuracies were comparable. We imputed most of the 42 high-coverage genomes downsampled to 1x with low error rates (below 5%) and estimated higher error rates for African genomes, which are underrepresented in the reference panel. We used the ancient trio data to validate imputation and phasing results using an orthogonal approach based on Mendel’s rules of inheritance. This resulted in imputation and switch error rates of 1.9% and 2.0%, respectively, for 1x genomes. We further compared the results of downstream analyses between imputed and high-coverage genomes, notably principal component analysis (PCA), genetic clustering, and runs of homozygosity (ROH). For these three approaches, we observed similar results between imputed and high-coverage genomes using depths of coverage of at least 0.5x, except for African genomes, for which the decreased imputation accuracy impacted ROH estimates. Altogether, these results suggest that, for most populations and depths of coverage as low as 0.5x, imputation is a reliable method with potential to expand and improve ancient DNA studies.

## Introduction

Ancient DNA (aDNA) is characterized by pervasive *postmortem* damage, including fragmentation and deamination^1^. As a result, most ancient genomes have low breadth and depth of coverage, hindering confident genotype calling. Instead, pseudo-haploid data are commonly generated by sampling one allele per variant site^2,3^. Evermore methods and tools are developed to study different aspects of population structure, including diploid genetic properties such as runs of homozygosity (ROH)^4^, using pseudo-haploid data. However, on the one hand, methods designed for diploid and haplotypic data cannot be easily applied to pseudo-haploid data, and, on the other hand, these data come with increased bias towards the reference genome^5^.

One alternative to downsampling the data to pseudo haploid prior to downstream analyses is to impute low-coverage ancient genomes. The goal of imputation is to infer missing sites, usually by using reference panels of haplotypes. Most imputation tools employ a hidden Markov model (HMM) that determines which assembly of reference haplotype chunks represents the target best. Mostly, the Li and Stephen model of linkage disequilibrium (LD)^6^ is at the core of this HMM. This model describes LD in terms of the subjacent recombination rates. In particular, it estimates the probability of observing a chromosome (or haplotype) given the already sampled haplotypes from a population by considering the new haplotype as a copy of different parts of the sampled haplotypes while allowing mutations to arise. The transition rate between copying haplotypes is proportional to the recombination rate and it decreases with the number of available haplotypes to copy from.

SNP-array imputation is applied when genomes are genotyped at a subset of variant sites^7^. SNP-array imputation of modern DNA is often implemented to increase required sample sizes for genome wide association studies (GWAS), so as to avoid the still high whole-genome sequencing (WGS) costs^8^. It is also possible to impute low-coverage genomes whose genotypes cannot be directly determined with certainty, in which case genotype uncertainty is captured by likelihoods, instead of hard calls^9–14^. One can make use of this second type of methods to impute low-coverage ancient genomes. Present-day genotypes have been imputed with increasing accuracy due to improved imputation methods on the one hand, and increased reference panel size and diversity on the other hand, such as the Haplotype Reference Consortium (HRC)^15^, the 1000 Genomes Project^16^ and TOPMed^17^. These advances have also been exploited by some (e.g., Martiniano et al., 2017^18^; Haber et al., 2020^19^; Saupe et al., 2021^20^; Clemente et al., 2021^21^, Cox et al., 2022^22^; Allentoft et al., 2022^23^) to impute low-coverage ancient genomes, using present-day haplotypes, assuming matching ancestry.

However, aDNA introduces extra challenges, including damage and potential contamination^24^, and it is not clear whether ancient individuals’ ancestries are well captured by reference panels of present-day individuals. Moreover, a precise quantification of possible imputation biases and errors is lacking. Hui et al.^25^ proposed a two-step imputation pipeline to be applied to ancient genomes. This pipeline first imputes based on genotype likelihoods using Beagle4.1^10^, and then removes sites based on their maximum genotype probability (GP), a measure of how likely each possible genotype at a site is to be true after imputation. The resulting genotype calls are again imputed with Beagle5^26^, followed by a final GP filtering step. When compared to the first imputation step alone, this pipeline yielded larger proportions of heterozygous sites that pass the specified GP threshold. Nonetheless, a single downsampled ancient European genome was used to validate these results. Another recent study^27^ assessed the imputation of ancient genomes performance by downsampling (0.1-2.0x) and imputing genomes from five high-coverage ancient Europeans using Beagle4.0^28^ and various reference panel and sample size configurations. The authors measured genotype concordance, bias towards the reference panel and compared projections of the high-coverage, low-coverage and imputed 1x data onto principal component analysis (PCA) of present-day data. Imputation accuracy improved when i) using all populations in the 1000 Genomes reference panel instead of restricting to European genotypes alone and ii) the ancient genomes were imputed simultaneously. They found no bias increase towards the most common reference panel allele for ancient genome coverages as low as 0.75x.

These two studies^25,27^ suggest that aDNA imputation performs well under specific conditions. However, in their assessment of imputation accuracy they used a rather limited sample of ancient genomes (one^25^ and five^27^) and of only European descent. Furthermore, more accurate and efficient low-coverage imputation methods are available, e.g., GLIMPSE^12^, than the methods they tested, i.e., Beagle4.0 and 4.1. Here, we make use of 43 ancient genomes, including an ancient trio and 42 high-coverage (>10x) genomes, from four different continents and different time spans to assess i) imputation accuracy of low-coverage ancient genomes and ii) how imputation affects downstream analyses. To this end, we downsampled to low coverage this diverse dataset of ancient genomes, which allowed us to quantify imputation performance across different ancestries, unlike, to our knowledge, any other previous study. We imputed the downsampled ancient genomes with GLIMPSE^12^, a state-of-the-art imputation and phasing tool that was shown to accurately impute low-coverage present-day genomes, having 1000 Genomes^16^ as a reference panel. In the next sections, we show how imputation accuracy varies with depth of coverage, substitution type, i.e., transitions vs. transversions, imputation methods, ancestry, and post-imputation filtering. To address our second goal, we assess the effects of imputation not only on PCA, but also on genetic clustering and ROH analyses.

## Results

The approach we followed in this study is schematically described in **Figure 1A**. We generated two datasets: imputed genotypes from downsampled genomes and corresponding validation genotypes called from the high-coverage ancient genomes, that we used as the ground truth. We started by sampling fractions of the sequencing reads from the 43 ancient genomes to obtain genomes with average depths of coverage between 0.1x and 2.0x. Then, using bcftools^29^ (on the choice of genotype caller prior to imputation in **Supplementary Section 1**), we generated genotype likelihoods at biallelic sites of the 1000 Genomes phase 3 v5 data^16^ phased with TOPMed^17^, the imputation reference panel, including all transition sites, in contrast to other studies^27^. We then imputed the data with GLIMPSE with the different steps described in the methods section. Lastly, we called genotypes for the high-coverage genomes and filtered out low-quality calls (methods and **Supplementary Section 2**), thus reducing the deamination impact. Finally, we assessed imputation performance and compared the downstream analyses results obtained with high-coverage and imputed genotypes.

**Figure 1:**
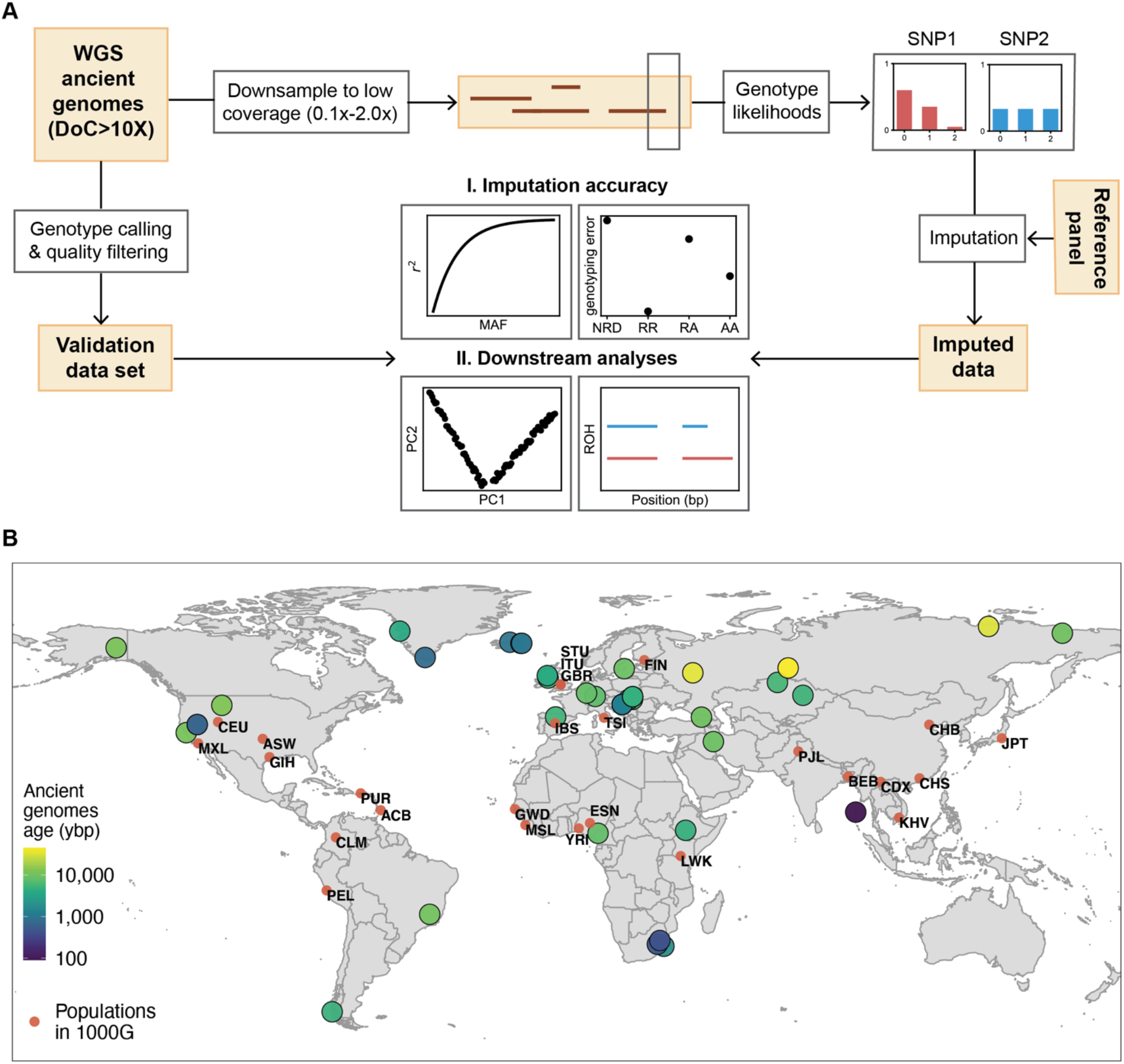
Overview of the procedure we followed (A), and geographical origin and age in years before present (ybp) of the 43 individual samples used in this study as well as the different populations represented in the 1000 Genomes reference panel (B).

Three out of the 43 ancient genomes in this study constitute a trio (mother, father and son) that was recently re-sequenced and is not yet fully public^23,30^, in contrast to the remaining 40 genomes. This dataset of 43 ancient genomes is a diverse dataset in regard to their sequencing/study, as well as epoch and continent the ancient individuals lived in, with about half of the individuals being from Europe and the other half from Africa, America and Asia (**Figure 1B**). Information concerning location and age of remains, and genome coverage is included in **Table S1**.

### 1. Accuracy of low-coverage ancient DNA imputation

We started by examining how imputation quality changes with average depth of coverage, and whether transversions are imputed more accurately than transitions, since the latter are affected by *postmortem* DNA deamination, i.e., C-to-T substitutions, which might wrongly increase the number of called heterozygous sites. We further compared imputation performance using two different state-of-the-art imputation methods, GLIMPSE and Beagle4.1^10^, where the latter is a widely used imputation method and was applied in Hui et al^25^. For that, we calculated imputation accuracy, r^2^, that is, the squared Pearson correlation between genotype dosage in the aggregate of the 42 high-coverage and imputed datasets, as a function of minor allele frequency (MAF) as determined from the 1000 Genomes reference panel.

#### Ancient and present-day DNA imputation accuracies are comparable

We found that imputation accuracy of ancient genomes was similar to the accuracy reported for present-day genomes when using the same imputation method^12^. Accuracy was higher for common variants (MAF≥5%) (**Figure 2A**), as rare variants are more challenging to impute^8,31^. Imputation accuracy was also higher for genomes with higher coverage, as these have more data. In particular, for depths equal and greater than 0.75x, we obtained r^2^>0.90 at sites with MAF>2%, and r^2^>0.70 and r^2^>0.95 for rare (0.1%<MAF≤1%) and common variants (MAF≥10%), respectively. We then found that GLIMPSE outperformed Beagle4.1 for 1x ancient genomes, particularly at rare variants (**Figure S3**), similarly to the case of present-day genomes^12^. Finally, we did not observe substantial differences in accuracy between imputed transversion and transition sites (**Figure S3**).

**Figure 2:**
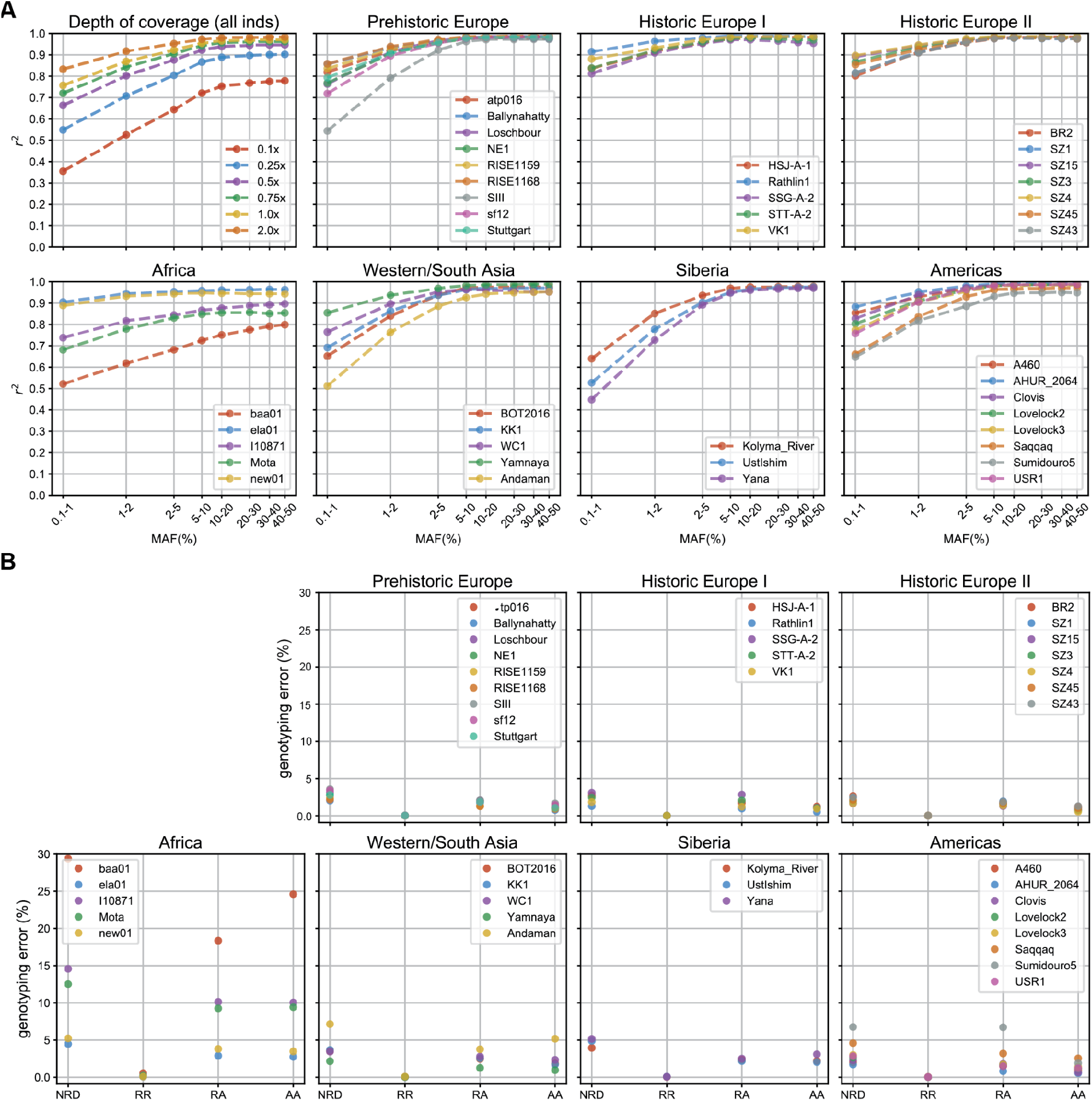
Imputation quality assessment: A) imputation accuracy (r^2^) as a function of minor allele frequency (MAF) for the 42 high-coverage genomes together downsampled to different depths of coverage (top left) and for individual 1x genomes (remaining plots); B) genotype discordance between individual imputed (1x) and high-coverage genomes for homozygous reference allele (RR), heterozygous (RA) and homozygous alternative allele (AA) sites, as well as the resulting non-reference discordance. Depending on ancestry, MAF was specified from the reference populations expected to be closer to the individual in question, whenever possible, as listed in **Table S1**. Individuals were put in categories that roughly reflect their place of origin and/or time.

Fixing depth of coverage at 1x, we evaluated how imputation performs across the 42 high-coverage genomes of different ancestries and times. In addition to imputation accuracy as a function of MAF, we quantified genotyping error rates for homozygous reference and alternative allele and heterozygous sites. We also report the non-reference discordance (NRD), that is, the ratio of the number of incorrectly imputed sites and the total number of imputed sites, excluding correctly imputed homozygous reference allele sites.

#### Imputation error rates below 5% for most non-African 1x genomes

The imputation of European, Western, and most Native American genomes yielded similar accuracy curves starting with lower values for rare variants (0.5<r^2^≳0.9) and converging to r^2^>0.90 from MAF≥2% (**Figure 2A**). The African ancient genomes were the least accurately imputed with only two out of five imputed genomes reaching r^2^>0.90, and error rates as high as 18% at heterozygous sites, the most challenging to impute, and NRD between 4% and 29% (**Figure 2B**). In contrast, most non-African imputed genomes yielded NRD rates below 5%. This difference in imputation performance is likely due to lack of representation of the different African populations in the reference panel. Although the 1000 Genomes reference panel contains individuals of African origin, mostly from West Africa (Mende Sierra Leone (MSL), Gambian Mandinka (GWD), Esan Nigeria (ESN), Yoruba (YRI) and Luhya Kenya (LWK)), the genetic diversity in Africa^32^ is not well represented in this panel. Therefore, reference populations from West Africa might not represent populations in Southern Africa^33^ for imputation purposes, as in the case of baa01^34^, the most poorly imputed ancient genome. Conversely, European ancient individuals are better represented in the reference panel. And yet, Native American genomes were also accurately imputed, even though the populations in the reference panel show different admixture moieties, ranging from low (e.g., Puerto Rican (PUR)) to high Native American (e.g., Peruvian (PEL))^16^ admixture proportions. This suggests that having haplotypes in the reference panel that match the ancestry of the target haplotypes is fundamental to achieve high imputation accuracy, even if these reference haplotypes originate from admixed individuals.

#### Validating imputation and phasing accuracy on an ancient trio

The availability of an ancient trio (mother, father, son) allowed us to use an orthogonal approach based on Mendel’s rules of inheritance to measure imputation and phasing quality. This trio was sampled in a Late Neolithic mass burial at Koszyce^23,30^ and was re-sequenced in the context of the study of Allentoft et al.^23^ resulting in genome coverages of 27.5x (mother, RISE1159), 18.9x (father, RISE1168), 5.4x (son, RISE1160). In this analysis, imputation errors corresponded to sites where parental and offspring genotypes disagreed with Mendel transmission rules. Here, we excluded sites that are homozygous for the reference allele in the three genomes as these positions are easier to impute. We estimated phasing accuracy in terms of switch error rate, that is assessed for every two consecutive heterozygous sites by verifying if the alleles for the two sites are located on the correct haplotypes following the expected configuration from the trio. Mendel error rates ranged from 1.3% at 4x to 12.2% at 0.1x (**Figure 3A**). For 1x data, in particular, Mendel error rates were between 1.5% and 2.9% across the 22 autosomes. These error rates agree with previously estimated imputation errors (**Figure 2B**). Switch error rates varied between 1.6% at 4.0x and 8.2% at 0.1x, with errors for 1x data in the range 1.6%-3.0% (**Figure 3B**). For present-day genomes and small sample sizes, switch error rates are typically between 1% and 5%^35–37^, and we achieved similar accuracy when imputing and phasing the genomes downsampled to a minimum coverage of 0.25x.

**Figure 3:**
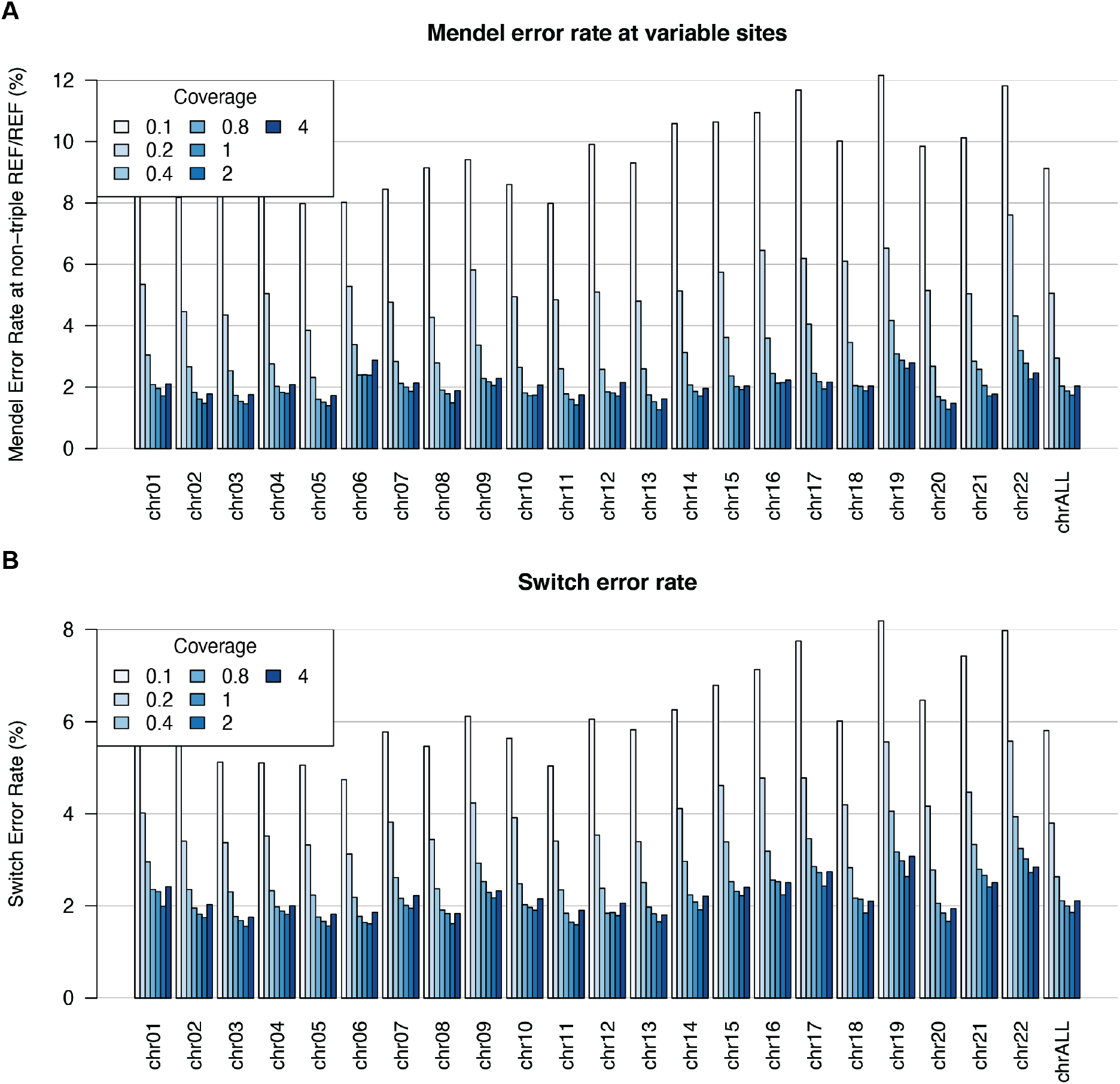
Imputation and phasing accuracy for the Koszyce trio: A) Mendel error rate across the 22 autosomes is counted when the parental and offspring genotypes violate Mendel transmission rules, excluding sites at which all three non-imputed genomes are REF/REF; B) switch error rates averaged over the three genomes. A switch error is counted between two consecutive heterozygous genotypes when the reported haplotypes are not consistent with those derived from the trio.

#### Genotype probability filtering: a trade-off between more accurate calls and alternative allele sites loss

After imputation, we can filter based on the maximum of genotype probabilities (GP) for a site. GP is a measure of how likely each genotype is to be true and takes values between 0 and 1 that sum to 1 across the possible genotypes. To determine which GP value we would use to filter the imputed data prior to downstream analyses, we applied GP filters starting at 0.70 and up to 0.99 to four different imputed ancient genomes downsampled to 0.1x and 1.0x (RISE1 168^23,30^, SIII^38^, Ust’-Ishim^39^ and Mota^40^). We then quantified imputation accuracy and genotype discordance. We observed a greater boost in accuracy as the GP filter becomes stricter for 0.1x imputed data than for 1x data (**Figure 4A**). In the case of 1x data, we obtained small improvements in accuracy for sites with MAF>5%. The exception was the individual sample Mota, where the gain in accuracy for a specific GP filter had similar magnitude across sites with different MAF values. This African genome yielded the second lowest imputation accuracy amongst the 42 ancient high-coverage genomes downsampled and imputed in this study. We observed the same trends with genotype discordance between imputed and high-coverage genotypes (**Figure 4B**). Genotyping error rates were higher for 0.1x than for 1x imputed genomes, for whom error rates remained below 5%, except for Mota. Increasing GP filtering values decreased these error rates in all instances. Then, we looked at how GP filtering affects the number of correctly imputed heterozygous sites (**Figure 4C**). The proportion of lost heterozygous sites was much higher in the case of 0.1x data, explained by the lower imputation accuracy for this coverage. For 0.1x data, filtering out sites with GP<0.70 removed around 15% of correct heterozygous sites in the least. When GP≥0.99, only between 20% and 43%of correct heterozygous sites remained. In contrast, the imputed 1.0x genomes lost a small fraction of their heterozygous sites as stricter GP filters were applied. This fraction was smallest amongst the genomes of European ancestry (<8%, RISE1168 and SIII) and largest for Mota (22%), a reflection of how accurately these genomes were imputed. In the end, a trade-off must be made between loss of heterozygous sites and imputation accuracy. Based on these results, we chose to remove sites with MAF<5% and set to missing imputed sites with GP<0.80, for most of the downstream analyses, thus keeping most heterozygous sites for 0.1x data while controlling for imputation accuracy.

**Figure 4:**
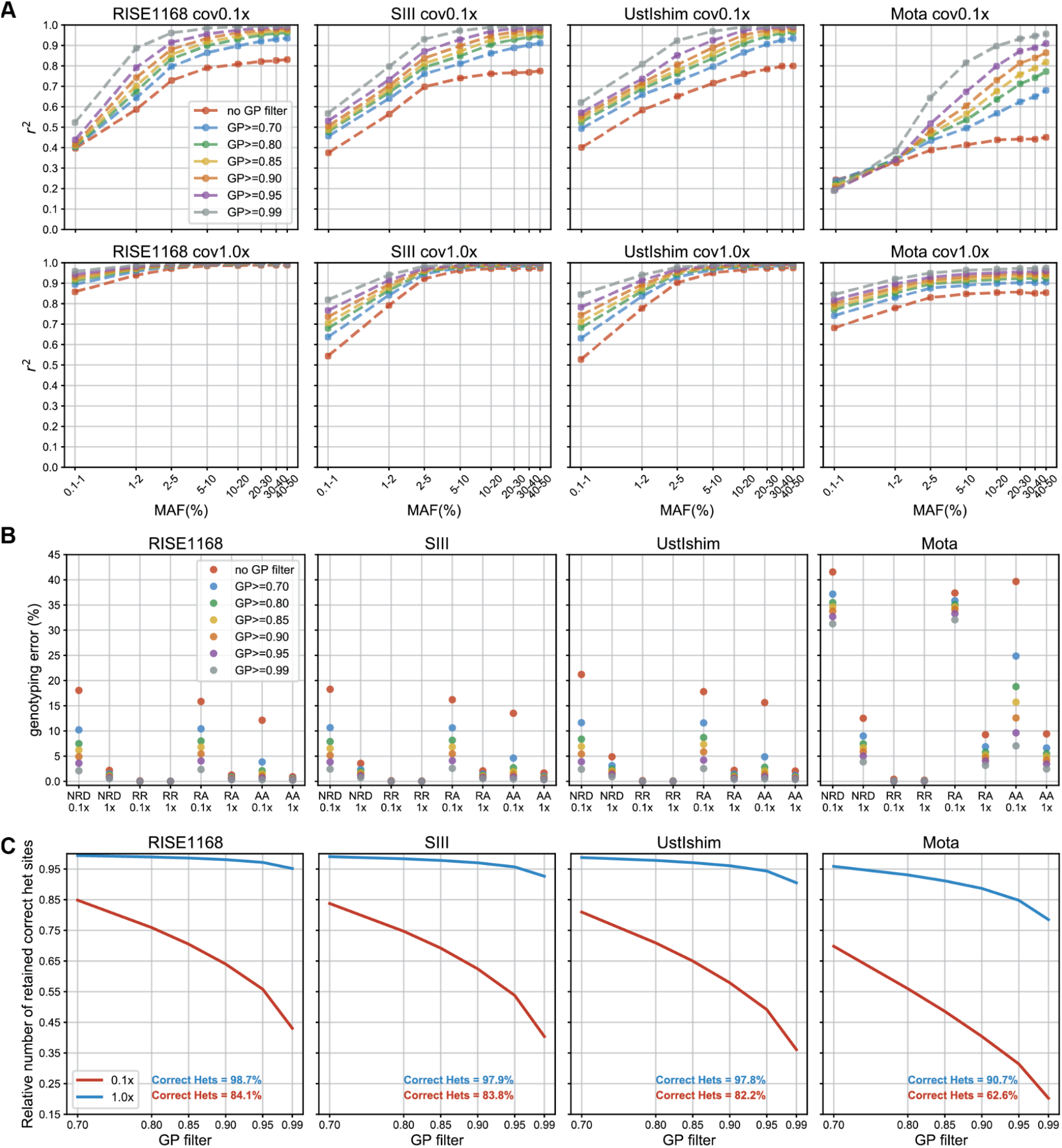
Effects of applying different thresholds when filtering for GP in the case of four imputed 1x ancient genomes (RISE1 168^23,30^, SIII^38^, Ust’-Ishim^39^ and Mota^40^) on A) imputation accuracy, B) genotype discordance between imputed and non-imputed genomes for homozygous reference allele (RR), heterozygous (RA) and homozygous alternative allele (AA) sites, and also the non-reference discordance (NRD), C) proportion of correctly imputed heterozygous sites retained for 0.1x and 1.0x data for each of the four genomes. The percentage of correctly imputed heterozygous sites for 0.1x and 1.0x before GP filtering are represented in red and blue, respectively, in panel C.

### 2. Imputation effect on downstream analyses

In order to detect and quantify potential bias introduced by imputation, we compared the results of downstream analyses, namely, principal component analysis (PCA) and genetic clustering analyses, performed with the high-coverage and imputed genomes, after filtering for MAF and GP (imputed data). These three methods are broadly used in population genetics to investigate population structure and demography. PCA is a dimension reduction technique that helps visualizing patterns of population structure. In the genetic clustering analyses, the ancestry of an individual is estimated as the sum of K different clusters determined from the data in an unsupervised fashion. We further explore the potential of imputing low-coverage ancient genomes by estimating runs of homozygosity (ROH), whose classical applications require diploid data. ROH segments are unbroken homozygous regions of the genome that contain information about past and recent breeding patterns^41^. ROH have been found in all populations, but their number and size vary, depending on demographic histories.

For the PCA, we calculated the first ten principal components of the 1000G reference panel and projected both the high-coverage and corresponding imputed ancient genomes onto those. We have included both transition and transversion sites in this analysis.

#### Imputation did not introduce significant bias in PCA for coverages of at least 0.5x

Both the imputed 1x and high-coverage ancient genomes were in the expected continental groups as defined by present-day individuals in the two first principal components (**Figure 5A**). They also tended to colocalize, which was particularly the case for ancient individuals clustering with present-day Europeans, suggesting limited bias is introduced by imputation in the PCA results. To further verify whether imputation introduced bias in this analysis, we took the difference in coordinates between validation and corresponding imputed 1x genomes for each principal component. As shown in **Figure 5B**, the normalized differences between the two datasets were small and did not deviate significantly from 0 (t-test p-values > 0.01). Additionally, we found that only genomes with coverage as low as 0.1x and 0.25x show some significant deviation from 0 (**Figure 5C**) for some principal components, however, the imputed data were still placed in the expected continental clusters in the PCA space (**Figure S4**). This is particularly clear for European ancient genomes. These results show that the differences between imputed and high-coverage coordinates tended to be centered on 0 for the first principal components, in particular for genomes with coverage above 0.25x, suggesting that imputation did not introduce a significant bias to the PCA.

**Figure 5:**
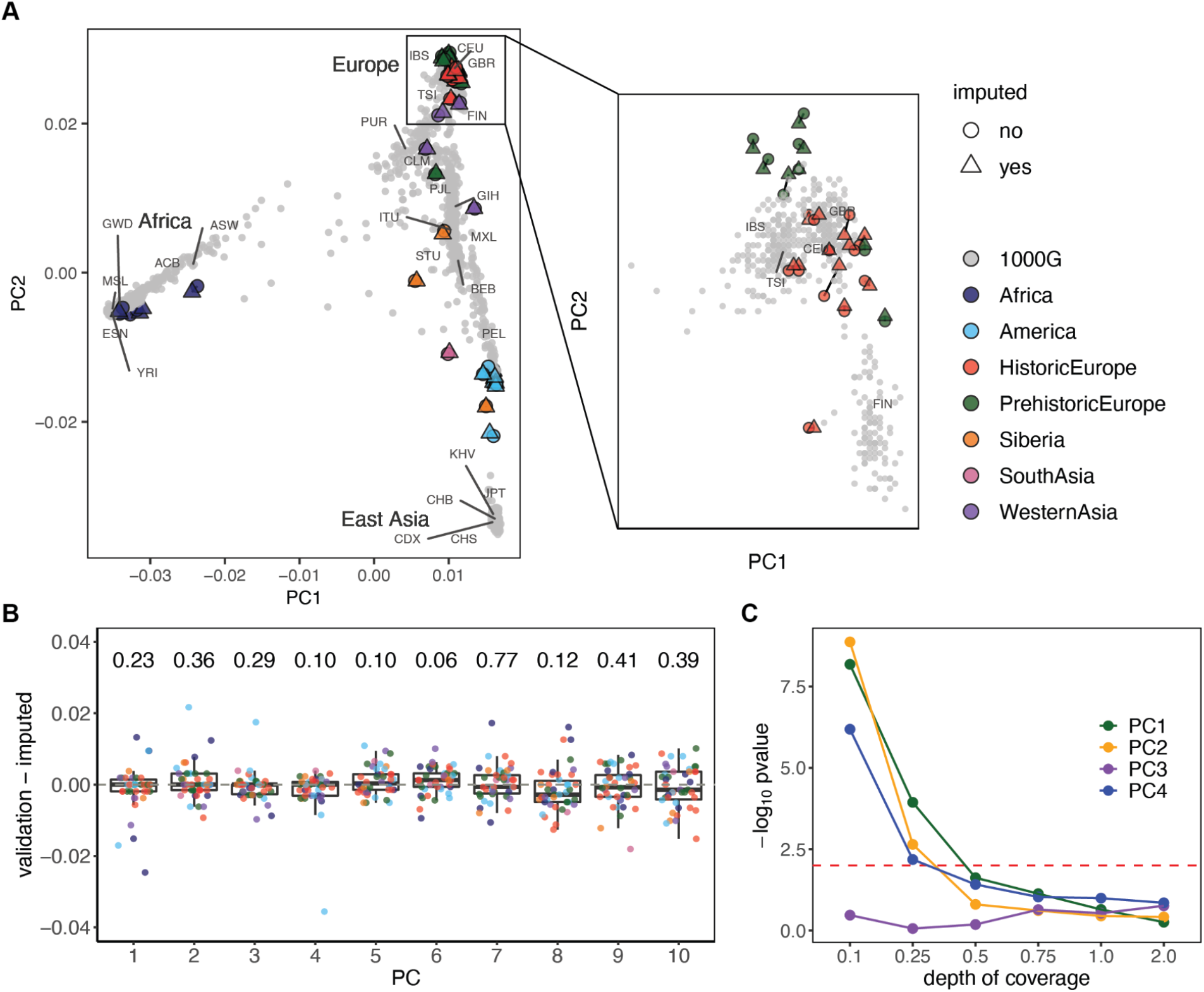
Principal component analysis (PCA) of imputed and high-coverage ancient genetic data, and present-day data in 1000 Genomes reference panel: A) projections for 1x imputed, high-coverage and present-day data along the first two principal components, where 1000 Genomes individuals are plotted in gray and population labels are shown in the average location of the individuals from the same population, ancient individuals are colored by region and/or epoch, with the high-coverage and imputed individuals represented by full circles and triangles, respectively; the plot on the left contains the coordinates of the whole data set and the plot on the right shows the coordinates of European modern individuals as well as of the European-labeled ancient individuals that cluster with these; B) boxplots of the normalized differences in coordinates between validation and corresponding 1x imputed genomes for the first 10 principal components and resulting p-values from testing whether differences are significantly different from 0; individual data points are overlaid and colored according to the region and/or epoch as in the previous plot; C) −log_10_ p-values obtained when testing whether differences between imputed and validation data are significantly different across the six depths of coverage and for the first four principal components; the red dashed line indicates a p-value of 0.01.

#### No ancestry bias in genetic clustering analyses of imputed European (≥0.5x) genomes

For the genetic clustering analyses, we focused on the European genomes. It is well established that the genetic diversity of present-day Europeans can be modeled with three ancestral populations: western hunter-gatherers, early European farmers and Steppe pastoralists^42^. Ancient European individual samples tend to exhibit different distributions of these three ancestries across time and space. We asked whether imputation of European ancient genomes artificially increases the amount of inferred Steppe-like ancestry for these individuals, since most present-day European individuals have Steppe ancestry, including the European populations in the 1000 Genomes reference panel. For instance, we assessed whether the Steppe-like component increases in imputed western hunter-gatherer genomes like Loshbour^42^. To this aim, we performed unsupervised admixture analyses with the software ADMIXTURE^43^, including transitions and transversions. We used as a reference panel the genetic data of 61 ancient individuals present in the 1240K dataset^44^, including nine western hunter-gatherers, 26 Anatolian farmers and 26 individuals of Steppe ancestry (see **Table S2**). We estimated ancestry proportions for the imputed and validation data separately varying the number of clusters (K) between two and five. For K=2, 4 and 5, we observe qualitatively similar results for imputed and high-coverage data (see **Supplementary Section 6**). Here we show the results obtained with K=3 (**Figure 6A**), as these clusters seemingly capture the three aforementioned ancestries. The admixture proportions are qualitatively similar between the high-coverage ancient genomes and the corresponding imputed ones, and, in the particular case of Loschbour, the only western hunter-gatherer imputed in this study, we estimated 100% western hunter-gatherer-like ancestry with both imputed 1x and high-coverage data (**Figure 6B**). In order to compare the admixture results across imputed data with different depths of coverage, we took the difference between ancestry proportions estimated for the validation and imputed genomes for each ancestry component and each coverage (**Figure 6C**). We observed larger differences with imputed 0.1x and 0.25x data. For the remaining depths of coverage, the small differences distributed around 0 show no indication that imputation introduced any substantial bias towards a particular ancestry in this analysis.

**Figure 6:**
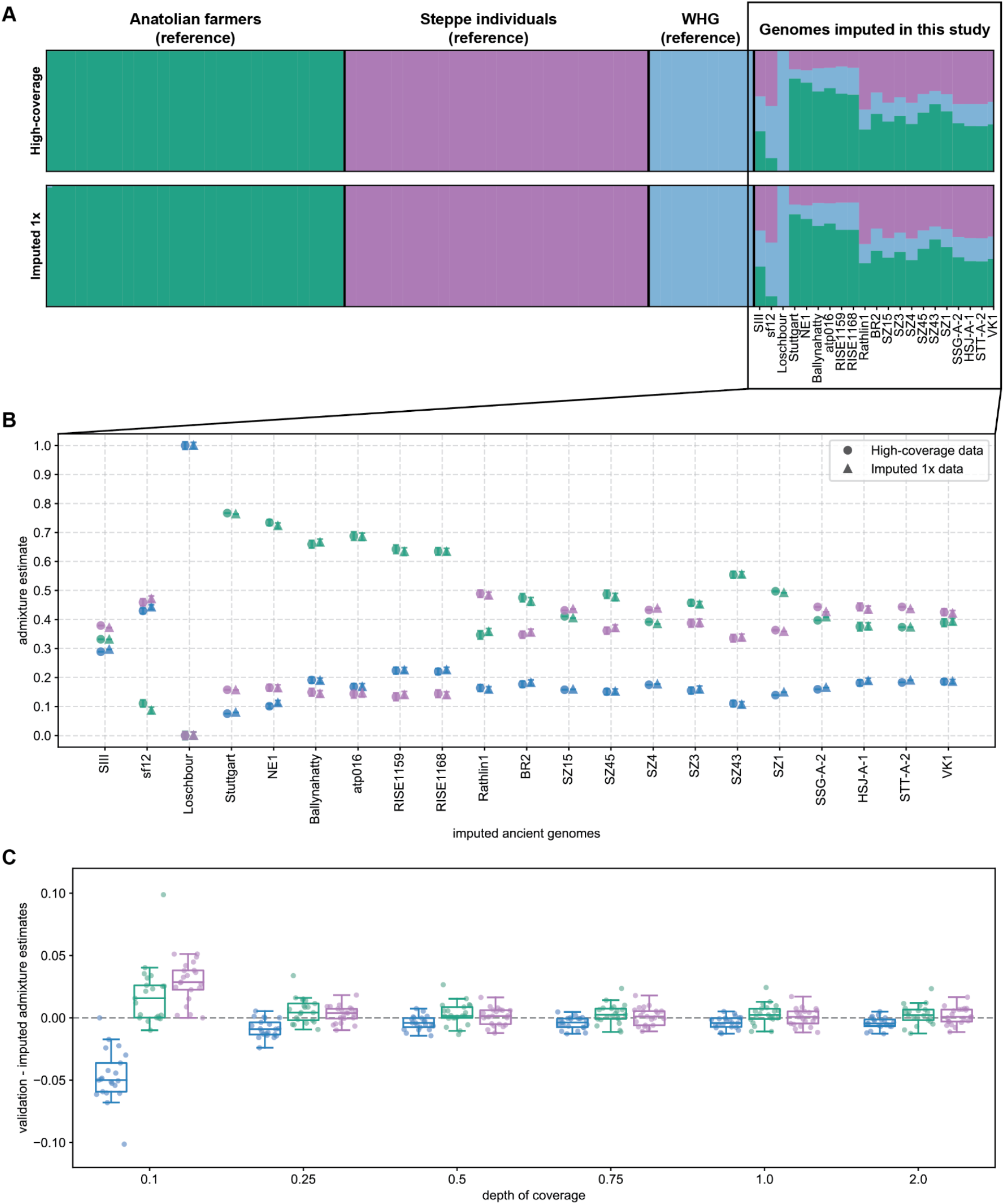
Unsupervised admixture analyses of European ancient individuals with three clustering populations: A) resulting admixture proportions and clusters for the reference and the 21 European individuals in this study, with validation results on top and imputed 1x below; B) admixture estimates for each of the three clusters obtained with imputed 1x (triangles) and validation (full circles) data for each of the 21 individuals, where error bars represent one standard error of the estimates; C) boxplots of the differences between the values of ancestry components obtained with the high-coverage and imputed data across all depths of coverage.

#### ROH estimated in imputed and high-coverage genomes overlap

Then, we first quantified ROH using transversions only to minimize the aDNA damage impact on the validation estimates. We examined how well the imputed and the validation ROH overlapped in chromosome 10 for each depth of coverage and for four different individuals, namely Ust’-Ishim^39^(Siberia), Rathlin1^45^ (Europe), A460^46^ (Americas) and Mota^40^ (Africa) (**Figure 7A**). The imputed 0.1x data had an excess of ROH when compared to the high-coverage data. This likely results from i) reduced imputation accuracy and ii) removal of a large proportion of heterozygous sites when applying post-imputation filters (**Figure 4C**). As the depth of coverage increased, the number of falsely identified ROH tended to decrease, while most validation ROH were also found amongst the imputation ROH. We then compared the total ROH lengths, stratified by segment size, measured in the imputed data with the validation data for the different depths of coverage and the same four individuals (**Figure 7B**). Again, we found the largest discrepancies between validation and imputed 0.1x data, with an excess of ROH segments, particularly of the shortest kind (0.5-1.0 Mb). For coverages above 0.1x, the total ROH lengths in the imputed genomes were close to the validation ROH. Lastly, restricting to imputed 1x data, we contrasted the total length of small ROH (<1.6 Mb) with the total length of longer ROH (⩾1.6 Mb) obtained with transversions only (**Figure 7C**) and all sites (**Figure 7D**). When using transversions only, the total ROH lengths estimated for high-coverage and corresponding imputed 1x genomes were similar, particularly for the European genomes. Furthermore, the ROH trends for the ancient individuals mostly agreed with documented ROH for their present-day counterparts, with Africans having the smallest total ROH lengths and Native Americans the longest^41^.

**Figure 7:**
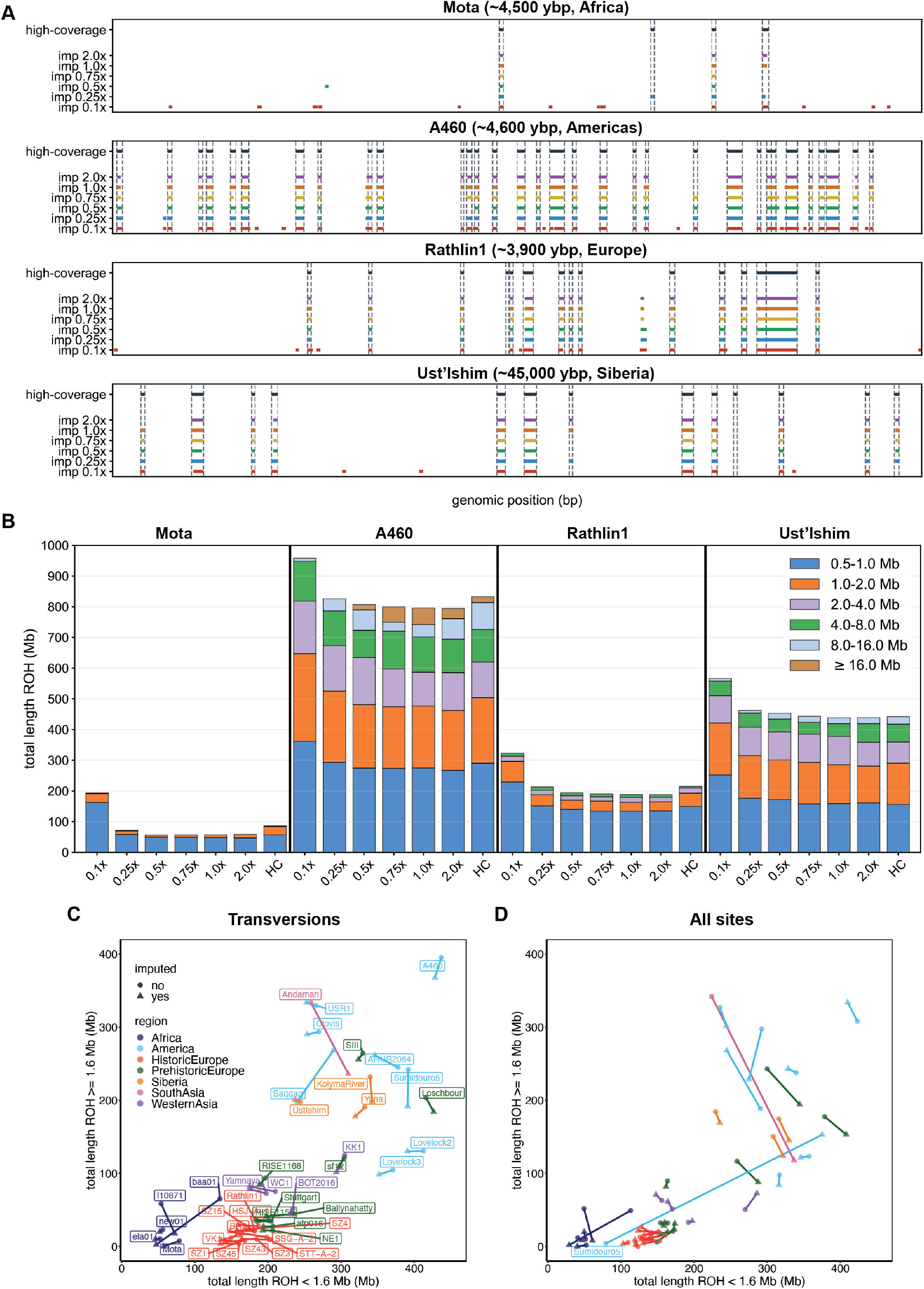
Runs of homozygosity (ROH) estimates for the high-coverage and corresponding imputed genomes: A) ROH locations in chromosome 10 found using transversions only with high-coverage and imputed genomes, in the case of four ancient individuals, namely, Mota^40^ (~4,500 ybp (years before present), Africa), A460^46^ (~4,600 ybp, Americas), Rathlin1^45^ (~3,900 ybp, Europe), Ust’-Ishim^39^ (~45,000 ybp, Siberia); B) total length of ROH discriminated by individual ROH length categories, estimated for imputed and high-coverage genomes (HC) using transversion sites for the four aforementioned individuals; C) total length of long (≥1.6 Mb) vs. small (<1.6 Mb) ROH segments for validation (full circles) and 1x imputed (triangles) genomes using transversion sites only and (D) using transversions and transitions.

#### Imputation seems to correct damage in ROH estimates in the case of Sumidouro5

When we added transitions to estimate ROH, the distance between imputed and validation ROH increased for some genomes (**Figure 7D**). In the case of the ancient Native American Sumidouro5^46^, this distance dramatically increased. The high-coverage estimate for Sumidouro5 was now located between the African and European values, but the imputed estimate remained close to both the high-coverage and imputed values obtained with transversions only. For this genome, we found major differences between high-coverage ROH sizes obtained with transversions only and all sites, whereas the corresponding imputed ROH were highly consistent (**Figure S12**). This indicates that the discordance between validation and imputed ROH, when transitions were introduced, originated from the validation data. Indeed, Sumidouro5 is a very damaged genome (40% deamination rate)^46^, which likely led to an excess of heterozygous calls in the high-coverage data, despite the quality filtering (see **Supplementary Section 2**).

## Discussion

Here we showed that low-coverage ancient genomes can be imputed with similar accuracy as modern genomes. In particular, we obtained accurate results at common variants, for coverages starting at 0.5x from MAF>5% (or at 0.75x from MAF>2%). However, this threshold is dependent on the ancient genomes’ ancestry. We observed that how well populations are represented in the reference panel can have a profound impact on imputation accuracy, with genotyping errors at alternative allele sites above 5% and up to 25% among African 1x genomes. These populations are underrepresented in the reference panel, whereas European genomes are better represented, and their imputation resulted in low error rates. Most Native American ancient genomes were also accurately imputed, and there are no reference populations with 100% Native American ancestry, but only with mixed ancestry. This result has far-reaching implications for the potential of imputing ancient genomes, since it is not guaranteed that there will be a present-day population that directly descends from the population which the ancient individual originates from without having admixed. Our results suggest that using admixed reference populations that share recent ancestry with the target ancient genomes can be enough in order to attain accurate imputation.

For most genomes, we obtained similar results with high-coverage and imputed data with coverages as low as 0.5x for the downstream analyses we carried out, i.e., PCA, admixture clustering and ROH estimation. Imputation did not introduce major bias for the first principal components, nor did it considerably increase the proportion of any of the three main ancestry components found in Europeans. The similarity of validation and imputed ROH segments is worthy of note, since ROH estimation typically requires reliable knowledge of genotypes, which is only available for high-coverage genomes. This means that ROH estimation methods designed for diploid data can become possible with low-coverage ancient genomes after imputation.

Although we did not remove transition sites prior to imputation, we found that transversion and transition sites were imputed with comparable accuracy. In fact, when we compared ROH estimates performed with transversions and all sites, we observed that imputation corrected ROH in the case of Sumidouro5, with 40% C-to-T mismatch frequency at the end of the reads. Given this observation, imputation of ancient genomes has the potential of correcting genotypes that are affected by damage and other sources of error. It remains to assess whether we can accurately impute contaminated ancient genomes in such a way that contaminating sequences do not contribute to the final genotypes.

We did not explore numerous genotype and haplotype-based applications that can greatly benefit from imputation of low-coverage ancient genomes, such as temporal selection scans and local ancestry inference. Moreover, genotype imputation, in general, is expected to improve as more and larger reference datasets become available. The recent release of 200K whole-genome sequences in the UK Biobank^47^, which can be used as a reference panel for imputation, offers an opportunity to improve imputation performance in the case of low-coverage European genomes, including ancient genomes, especially at rare variants and lower depths of coverage. In the case of ancient DNA, when the target genome is not well represented by modern reference populations or when a boost in imputation accuracy is required, additional reference panels can be assembled with high-quality ancient genomes of individuals with more closely shared ancestry. Furthermore, the number of sequenced ancient genomes has been growing exponentially and with no sign of slowing down. This means that more and more ancient genomes will be available with different ancestries and from different periods and with that comes the opportunity to expand existing reference panels with ancient genomes and to implement imputation in a more standardized way.

## Methods

In this section, we describe the methods implementation, starting with imputation, that includes all the file processing, imputation using GLIMPSE and using Beagle4.1, then the three downstream applications (PCA, genetic clustering analyses and ROH) and finishing with the two reference data sets used in this study.

### 1. Imputation

#### a. File processing prior to imputation

We downsampled high-coverage (10x-59x range) ancient genomes to coverages 0.1x, 0.25x, 0.5x, 0.75x,1.0x and 2.0x, using samtools^29^ v1.10. Then, we computed genotype likelihoods for the downsampled and the original high-coverage genomes for variant sites present in the 1000 Genomes phase 3 reference panel^16^ phased with TOPMed^17^ (see methods section 3.a).

To generate the genotype calls and genotype likelihoods, we used bcftools^29^ v1.10 and, as default, the command bcftools mpileup with parameters *-I -E -a ‘FORMAT/DP’ --ignore-RG,* followed by bcftools call *-Aim -C alleles.* To call genotypes from the high-coverage genomes, we have applied additional parameters for quality control (more details below).

We also generated both genotype calls from the high-coverage genomes and genotype likelihoods for the downsampled data (1x) with ATLAS^48^ v0.9.9 (see **Supplementary Section 1** and **Supplementary Section 2**) using the MLE caller and the empirical post-mortem damage (PMD) pattern observed across reads, as described in https://bitbucket.org/wegmannlab/atlas/wiki. For sake of time, we skipped the first step, splitMerge, that separates single-end alignments by length and merges the mates of paired-end reads and requires specification of the different libraries contained in a bam file. It is often the case that an ancient genome is obtained from a mixture of paired-end and single-end libraries. We observed that this first step we skipped did not have much impact when the bam files only had single-end libraries, but the genotype calling was seemingly less accurate when there were paired-end libraries in the bam files. So, we do not report here results we obtained from ATLAS calls from ancient genomes that were sequenced from paired-end libraries.

To obtain a trimmed validation dataset (see **Supplementary Section 2**), we trimmed five base pairs at both ends of the reads using the command trimBam from the package bamutil^49^ v1.0.14. Then, we called genotypes using bcftools v1.10, as previously described.

The final validation dataset was obtained by implementing the following filtering approach^46^: i) genotype calling with bcftools v1.10 with mapping and base quality filters of 30 and 20 *(-q 30 -Q 20),* respectively, and with the parameter *-C 50,* as recommended by the SAMtools developers for BWA mapped data to reduce mapping quality for reads with an excess of mismatches; ii) exclusion of the sites that are not in the 1000 Genomes accessible genome strict mask^50^; iii) removal of sites located in regions known to contain repeats (RepeatMask regions in UCSC Table Browser^51^, http://genome.ucsc.edu/); iv) filtering out sites with extreme values of depth of coverage when comparing to the average genome coverage: below the maximum of one third of the mean depth of coverage *(DoC)* and eight, that is, 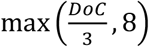, and depth above twice the average depth; v) filtering out of sites with the field QUAL below 30.

#### b. Imputation using GLIMPSE

We imputed the downsampled genomes using GLIMPSE^12^ v1.1.1. First, we used GLIMPSE_chunk to split chromosomes into chunks of sizes in the range 1 – 2 Mb and included a 200-kb buffer region at each side of a chunk. Second, imputation was performed with GLIMPSE_phase on the chunks with parameters *--burn* 10, *--main* 15 and *--pbwt-depth* 2, with 1000 Genomes as the reference panel. And then, we ligated the imputed chunks with GLIMPSE_ligate.

#### c. Imputation using Beagle4.1

To evaluate how GLIMPSE performs compared to Beagle4.1^10^ regarding imputation of low-coverage ancient genomes, we imputed the same data, but restricted to 1.0x, with Beagle4.1 with parameters *--modelscale* 2 and *--niterations* 0, that represent a trade-off between accurate results and running times.

#### d. Imputation accuracy evaluation

We used GLIMPSE_concordance to quantify imputation accuracy and genotype concordance, having the high-coverage data as validation. Only sites that were covered by at least eight reads and whose genotypes have a posterior probability of 0.9999 or more were used in validation. With GLIMPSE_concordance we obtained (i) imputation accuracy, that is, the squared correlation between dosage fields VCF/DS (DS varies between 0 and 2 that can be seen as a mean genotype value obtained from the genotype probabilities: 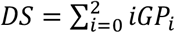, where *GP_i_*, is the genotype probability for genotype *i*) in imputed and validation datasets, divided in MAF bins, and (ii) genotype discordance, i.e., proportion of sites for which the most likely imputed genotype is different from the corresponding validation genotype for homozygous reference allele (RR), heterozygous (RA) and homozygous alternative allele sites (AA). We also estimated non-reference-discordance, NRD, defined as *NRD* = (*e_RR_* + *e_RA_* + *e_AA_)/(m_RA_* + *m_AA_* + *e_RR_* + *e_RA_* + *e_AA_*), where *e_X_* and *m_X_* stand for the number of errors and matches at sites of type X, respectively. NRD is an error rate which excludes the number of correctly imputed homozygous reference allele sites, which are the majority, thus giving more weight to imputation errors at alternative allele sites.

### 2. Downstream analyses

#### a. File processing

We filtered the imputed data by imposing that, for each variant site, the genotype probability (VCF/GP) for the most confidently imputed genotype to be at least 0.80. Then, we generated two datasets with different minor allele frequency (MAF) filters: MAF>5% (6,550,734 SNPs) for the data used in PCA and ROH analyses, and MAF>1% (11,553,877 SNPs) for admixture analysis, since with stricter MAF filters we would lose sites that distinguish the different populations. We used PLINK^52^ v1.90 to merge 1000 Genomes, high-coverage and imputed data into one file. In the case of PCA and admixture analyses, we intersected the resulting sites with the ones present in the Allen Ancient DNA Resource (AADR) data genotyped at the 1240K array sites^44^, that we refer to as the “1240K dataset” hereafter.

#### b. PCA

We performed PCA with smartpca (eigensoft^53^ package v7.2.1) without outlier removal (*outliermode: 2*). The 10 first principal components (*numoutevec: 10*) were calculated using the 1000 Genomes genetic data and both the imputed and high-coverage data were projected onto the resulting components (*lsqproject: YES*).

To perform the t-tests to test if there were significant differences in coordinates between validation and corresponding 1x imputed genomes for the first 10 principal components, we used the default R function *t.test*, running it in unpaired mode to test whether the mean of the differences was significantly different from 0 with a two-sided alternative hypothesis.

#### c. Admixture analysis

We estimated admixture proportions for 21 ancient Europeans with the software ADMIXTURE^43^ v1.3.0 in unsupervised mode. For the reference panel, we used a subset of the 1240K dataset containing nine western hunter gatherers, 26 Anatolian farmers and 26 individuals of Steppe ancestry^44^ (see **Table S2**). Contrary to the imputed and high-coverage genomes, the reference data are pseudo-haploid. We merged the reference panel with each of the imputation datasets (different coverages) with plink v1.90. We removed sites that were missing in more than 30% of the individuals. We proceeded similarly for the high-coverage dataset. We ran ADMIXTURE on seven configurations: merged reference panel and high-coverage individuals, and merged reference panel with each of the six imputed data sets (with initial coverage between 0.1x and 2.0x). For each configuration and number of clusters, we ran ADMIXTURE for K between two and five with 20 replicates (20 different seeds) and chose the replicate that yielded the largest log-likelihood value. In the final run, we obtained the standard error and bias of the admixture estimates using the option --*B 1000* that calculates these quantities with bootstrapping and 1000 replicates.

#### d. Runs of homozygosity (ROH)

We estimated ROH with plink v1.90 with the parameters^45^ --*homozyg*, --*homozyg-density* 50, --*homozyg-gap* 100, --*homozyg-kb* 500, --*homozyg-snp* 50, --*homozyg-window-het* 1, --*homozyg-window-snp* 50 and --*homozyg-window-threshold* 0.05. We estimated ROH twice: i) using transversion sites only, thus excluding sites that can be affected by aDNA damage, and ii) using both transversions and transitions.

### 3. Datasets

#### a. Ancient genomes in this study

The 43 downsampled and imputed ancient genomes (**Table S1**) were obtained from the “Ancient Genomes dataset” that was compiled in the context of the study of Allentoft et al^23^.

#### b. Reference panel for imputation

We used a version of 1000 Genomes v5 phase 3 (2,504 genomes)^16^, where the genomes were re-sequenced at 30x, and subsequently phased using TOPMed^17^, and with sites present in TOPMed. Only biallelic sites were retained (~90 million SNPs) and singletons were excluded. This panel was lifted over from build 38 to hg19 reference genome assembly using Picard liftoverVCF (https://gatk.broadinstitute.org/hc/en-us/articles/360037060932-LiftoverVcf-Picard-), with hg38ToHg19 chain from the University of California, Santa Cruz liftOver tool (http://hgdownload.cse.ucsc.edu/goldenpath/hg38/liftOver/).

#### c. Reference panel for genetic clustering analyses

We extracted a subset of the 1240K dataset^44^ containing ancient individuals of the three ancestries we were interested in: 26 Anatolian farmers (Anatolia_N), 26 Steppe individuals (Steppe_EMBA), and nine western-hunter gatherers (WHG), as specified in **Table S2**, to the exclusion of Loschbour, a genome that was also included in the dataset of 42 high-coverage genomes that we downsampled and imputed. We converted this subset from eigenstrat format to plink bed using the convertf command (eigensoft package v7.2.1). After that, we used plink v1.190 to do all of the data handling, such as merging plink bed files and filtering out sites with high missingness.

## Supporting information

Supplementary sections, including supplementary figures and tables

## Author contributions

B.S.d.M., A.-S.M. and O.D. designed the study and drafted the paper. B.S.d.M. and O.D. performed the experiments. S.R. helped with imputation. D.I.C.D., C.E.G.A, M.E.A., M.S. and E.W. helped with the population genetics analyses. H.S., M.E.A., N.N.J., M.H.S., P.W., A.S., M.M.P. generated and provided the ancient trio data. This work has been supervised by O.D. and A.-S.M. All authors helped with interpretation and reviewed the final manuscript.

## Acknowledgments

B.S.d.M. was supported by a Swiss National Science Foundation (SNSF) project grant (PP00P3_176977) to O.D. and by a European Research Council grant (grant agreement no. 679330) to A.-S.M. S.R. was supported by Swiss National Science Foundation (SNSF) project grant (PP00P3_176977). D.I.C.D. was supported by the European Research Council grant (grant agreement no. 679330) to A.-S.M. C.E.G.A. was supported by the National Institute of General Medical Sciences of the National Institutes of Health under award number R35GM142939. N.N.J. was supported by Aarhus University Research Foundation. H.S. was supported by the European Research Council (grant agreement no. 101045643).

