## Supplementary sections, including supplementary figures and tables for "Imputation of ancient genomes"

@ Joint last authors

### 1. Effect of using different genotype callers to generate genotype likelihoods, i.e., input for imputation

In order to determine whether the choice of genotype caller to calculate genotype likelihoods prior to imputation influences the quality of the imputed calls, we compared the accuracy of imputation using genotype likelihoods obtained with two different callers i) bcftools[1], a tool designed to handle modern DNA, and ii) ATLAS[2], a genotype caller that models and/or estimates deamination patterns and takes these into account when calling genotypes for ancient genomes. In the case of ATLAS, we empirically estimated the damage pattern before proceeding to genotype calling. In addition, to compute the imputation accuracy, we used two different validation sets, where calls were obtained using i) bcftools and ii) ATLAS. **Figure S1** shows imputation accuracy obtained with the four previously described configurations for a subset of 16 genomes downsampled to 1.0x prior to imputation. For a few genomes, such as Lovelock2[3] and SIII[4], there were no noticeable differences between the different configurations. In most cases, the most striking differences were between validation sets, regardless of the genotype likelihoods set used for imputation. Indeed, the accuracy curves tend to cluster by validation rather than genotype likelihood set. However, in the case of Sumidouro5, there is a larger difference between the two different genotype likelihood configurations for the same validation set, particularly at sites with minor allele frequency (MAF) below 5%. For this genome, with 40% frequency of C-to-T substitutions at the reads' ends, the highest accuracy was obtained with both imputation using genotype likelihoods and validation calls obtained with ATLAS. In conclusion, imputation calls were not significantly affected by the choice of tool to call genotype likelihoods for most of the cases here analyzed and, therefore, we chose to calculate genotype likelihoods using bcftools with no further filtering before imputation. However, the two genotype callers used to obtain the validation calls from the high-coverage genomes had clear differences and we further investigated the differences between them in the next section.

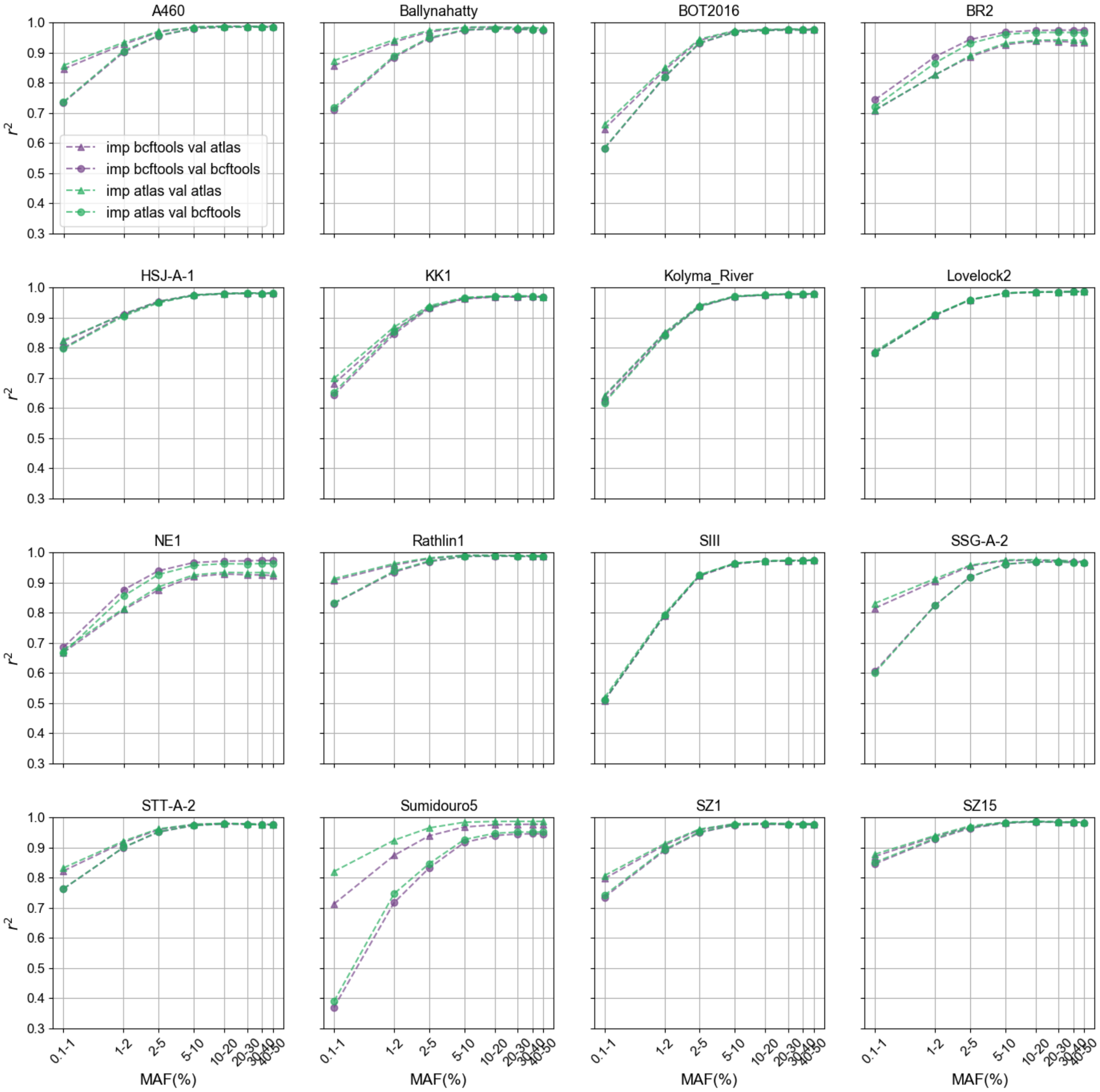

Figure S1: Imputation accuracy for a subset of 16 imputed 1.0x genomes, where imputation was performed from genotype likelihoods calculated with i) bcftools (purple) and ii) ATLAS (green). Two different validation sets were used in this analyses that differ in the tool used to call genotypes: i) bcftools (circles) and ii) ATLAS (triangles).

### 2. Validation dataset

In order to be able to assess imputation accuracy when imputing low-coverage ancient genomes, we resorted to downsampling and imputing ancient genomes with average depth of coverage above 10x, using the high-coverage genomes as ground truth. However, these high-coverage genomes are not free from the inherent ancient DNA challenges, in particular, base deamination. This constitutes a problem in determining what the true genotypes are for a particular genome, thus affecting how well we can validate imputation results in the context of this work. To determine how we can best circumvent this problem, we tried different approaches to generate the validation calls that were expected to decrease the impact of ancient DNA damage on the resulting genotype calls in the case of four 1x ancient genomes, namely, RISE1168[5,6], SIII[4] (UDG-treated), Sumidouro5[3] (a highly damaged genome) and WC1[7] (with intermediately high damage levels). Firstly, we called genotypes with ATLAS, a genotype caller that models the deamination patterns at the ends of reads, except for RISE1168, that includes paired-end libraries in addition to single-end libraries. Then, prior to genotype calling with bcftools, we trimmed five base pairs from the reads in the bam files. We also followed the filtering approach carried out in previous studies[3]: i) we called genotypes with bcftools using reads with mapping quality of at least 30 and bases with quality 20 ("-q 30 -Q 20"), as well as the recommended option -C 50; ii) we retained only the sites present in the 1000 Genomes accessible genome strict mask; iii) we excluded sites located in regions known to contain repetitions; iv) we removed sites that have depth below the maximum of one third of the mean depth of coverage and eight, as this is typically the minimum depth of coverage for which we can confidently call genotypes, and sites that have depth above twice the mean depth; v) we retained sites for which the field QUAL is equal or greater than 30. Finally, we used these different genotype call sets as validation when evaluating imputation accuracy for the four aforementioned genomes. We learnt that the application of the aforementioned five filters gives a consistently higher agreement between imputation and validation calls, even though it does not yield the highest accuracy for the four genomes (**Figure S2**). In the case of Sumidouro5, ATLAS outperformed the other approaches, with the five filters approach in second place, but ATLAS performance was not consistent across samples. The trimming approach yielded intermediate accuracy curves, in general, but, compared to the five-filter option in the case of Sumidouro5, it yielded much lower values at rare variants (MAF<2%). Given these results, we chose to generate the validation calls used in all the analyses in the main paper by applying the five-filter approach. However, we are aware that more accurate calls could be obtained by applying stricter filters for particular genomes that contain more degradation, including Sumidouro5, as was done in Moreno-Mayar et al.[3] We decided, instead, to apply the same approach to all 42 high-coverage genomes to expedite the process and have consistent datasets.

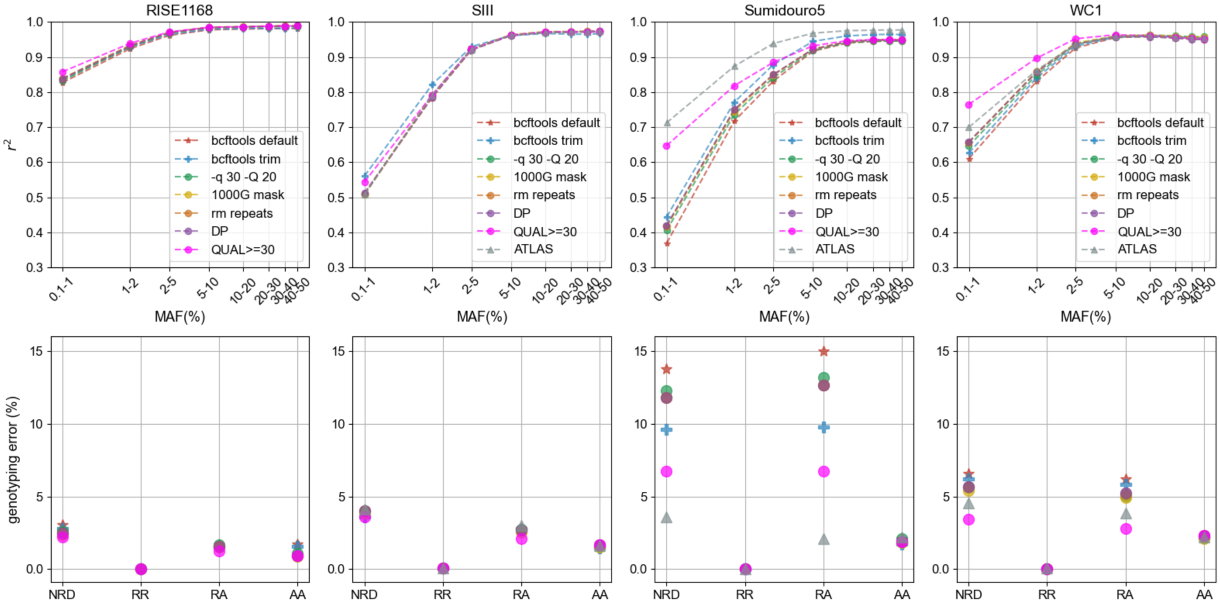

Figure S2: Effect of applying quality filters to validation call set on imputation accuracy (first row) and genotype discordance (second row) for four different genomes, organized by column (RISE1168[5,6], SIII[4], Sumidouro5[3] and WC1[7]). The imputed genomes had been downsampled to 1x coverage. The filters were applied in the following order: i) genotype calling with bcftools using reads with mapping quality of at least 30 and base quality 20 (“-q 30 -Q 20”), as well as the recommended option -C 50; ii) only sites present in the 1000 Genomes strict mask are retained; iii) exclusion of sites that are located in regions known to contain repetitions; iv) filtering out of sites that have depth below the maximum of one third of the mean depth of coverage and eight and sites that have depth above twice the mean depth; v) retaining sites for which the field QUAL is equal or greater than 30. For each validation and filter, the same imputed data were used in this analysis.

### 3. Individual samples

Table S1: Information on the ancient genomes used in the benchmark of imputation of low-coverage genomes: id, modern country where remains were found, age of remains in years before present (yBP), depth of coverage, and population of the 1000 Genomes panel whose minor allele frequency (MAF) was used in imputation accuracy analyses for each of the individuals, when applicable. AFR: Africa, AME: America, EAS: East Asia, EUR: Europe, SAS: South East Asia, All: overall allele frequency in 1000 Genomes. For RISE1160, a low-coverage genome, we do not indicate a MAF label, as we did not estimate imputation accuracy as a function of MAF for this genome. In the case of the ancient trio age (RISE1159, RISE1160 and RISE1168), we report a time span obtained for the mass grave as a whole. This range is the result of a model that took into consideration the number of contemporaneous individuals in the grave, radiocarbon dating of the different remains, as well as the ontogenetic constraints on how much the ages (e.g., parent/offspring relations) can vary[6].

| ID | Country | Age range (yBP) | Coverage | MAF label | Study |
|---|---|---|---|---|---|
| atp016 | Spain | 4867-5212 | 13 | EUR | Valdiosera et al., *PNAS* (2018)[8] |
| Stuttgart | Germany | 7020-7260 | 16 | EUR | Lazaridis et al., *Nature* (2014)[9] |
| Loschbour | Luxembourg | 7940-8160 | 18 | EUR | Lazaridis et al., *Nature* (2014)[9] |
| Ballynahatty | Ireland | 4970-5293 | 10 | EUR | Cassidy et al., *PNAS* (2016)[10] |
| sf12 | Sweden | 8757-9033 | 59 | EUR | Günther et al., *PloS Biology* (2018)[11] |
| NE1 | Hungary | 7021-7256 | 18 | EUR | Gamba et al., *Nat. Com.* (2014)[12] |
| RISE1159 | Poland | 4726-4830 | 27 | EUR | Schroeder et al., *PNAS* (2019)[6]; Allentoft et al., *bioRxiv* (2022)[5] |
| RISE1160 | Poland | 4726-4830 | 5 | - | Schroeder et al., *PNAS* (2019)[6]; Allentoft et al., *bioRxiv* (2022)[5] |
| RISE1168 | Poland | 4726-4830 | 19 | EUR | Schroeder et al., *PNAS* (2019)[6]; Allentoft et al., *bioRxiv* (2022)[5] |
| SIII | Russia | 33031-35154 | 11 | EUR | Sikora et al., *Science* (2017)[4] |
| Rathlin1 | Ireland | 3835 – 3976 | 11 | EUR | Cassidy et al., *PNAS* (2016)[10] |
| SSG-A-2 | Iceland | 950 – 1100 | 10 | EUR | Ebenesersdottir et al., *Science* (2018)[13] |
| HSJ-A-1 | Iceland | 950 – 1080 | 29 | EUR | Ebenesersdottir et al., *Science* (2018)[13] |
| STT-A-2 | Iceland | 950 – 1050 | 14 | EUR | Ebenesersdottir et al., *Science* (2018)[13] |
| VK1 | Greenland | 750 – 950 | 12 | EUR | Margaryan et al., *Nature* (2020)[14] |
| BR2 | Hungary | 3060 – 3220 | 18 | EUR | Gamba et al., *Nat. Com.* (2014)[12] |
| SZ15 | Hungary | 1346 – 1538 | 11 | EUR | Amorim et al., *Nat. Com.* (2018)[15] |
| SZ3 | Hungary | 1346 – 1538 | 11 | EUR | Amorim et al., *Nat. Com.* (2018)[15] |
| SZ4 | Hungary | 1347 – 1538 | 10 | EUR | Amorim et al., *Nat. Com.* (2018)[15] |
| SZ45 | Hungary | 1347 – 1538 | 10 | EUR | Amorim et al., *Nat. Com.* (2018)[15] |
| SZ43 | Hungary | 1347 – 1538 | 12 | EUR | Amorim et al., *Nat. Com.* (2018)[15] |
| SZ1 | Hungary | 3220 – 5320 | 11 | EUR | Amorim et al., *Nat. Com.* (2018)[15] |
| baa01 | South Africa | 1831 – 1986 | 14 | AFR | Schlebusch et al., *Science* (2017)[16] |
| ela01 | South Africa | 453 – 533 | 13 | AFR | Schlebusch et al., *Science* (2017)[16] |
| new01 | South Africa | 327 – 508 | 11 | AFR | Schlebusch et al., *Science* (2017)[16] |
| I10871 | Cameroon | 7800 – 7970 | 15 | AFR | Lipson et al., *Nature* (2020)[17] |
| Mota | Ethiopia | 4419 – 4525 | 10 | AFR | Gallego Llorente et al., *Science* (2015)[18] |
| KK1 | Georgia | 9550 – 9890 | 12 | EUR | Jones et al., *Nat. Com.*, (2015)[19] |
| WC1 | Iran | 9032 – 9405 | 10 | EUR | Broushaki et al., *Science* (2016)[7] |
| BOT2016 | Kazakhstan | 5318 – 5582 | 14 | EUR | Damgaard et al., *Science* (2018)[20] |
| Yamnaya | Kazakhstan | 4837 – 4968 | 26 | EUR | Damgaard et al., *Science* (2018)[20] |
| Andaman | India | 30 – 150 | 17 | SAS | Moreno-Mayar et al., *Science* (2018)[3] |
| UstIshim | Russia | 42560 – 47480 | 35 | All | Fu et al., *Nature* (2014)[21] |
| Yana | Russia | 30950 – 32950 | 27 | All | Sikora et al., *Nature* (2019)[22] |
| Kolyma_River | Russia | 9665 – 9906 | 15 | All | Sikora et al., *Nature* (2019)[22] |
| USR1 | USA | 11270 – 11600 | 17 | AME | Moreno-Mayar et al., *Nature* (2018)[3] |
| AHUR_2064 | USA | 10770 – 11170 | 19 | AME | Moreno-Mayar et al., *Science* (2018)[3] |
| Lovelock2 | USA | 1818 – 1942 | 15 | AME | Moreno-Mayar et al., *Science* (2018)[3] |
| Lovelock3 | USA | 567 – 687 | 19 | AME | Moreno-Mayar et al., *Science* (2018)[3] |
| Saqqaq | Greenland | 3600 – 4170 | 13 | AME | Rasmussen et al., *Nature* (2010)[23] |
| Clovis | USA | 12572 – 12726 | 15 | AME | Moreno-Mayar et al., *Science* (2018)[3] |
| Sumidouro5 | Brazil | 10258 – 10552 | 16 | AME | Moreno-Mayar et al., *Science* (2018)[3] |
| A460 | Chile | 4430 – 4850 | 11 | AME | Moreno-Mayar et al., *Science* (2018)[3] |

### 4. Imputation accuracy for transitions and transversions and method comparison

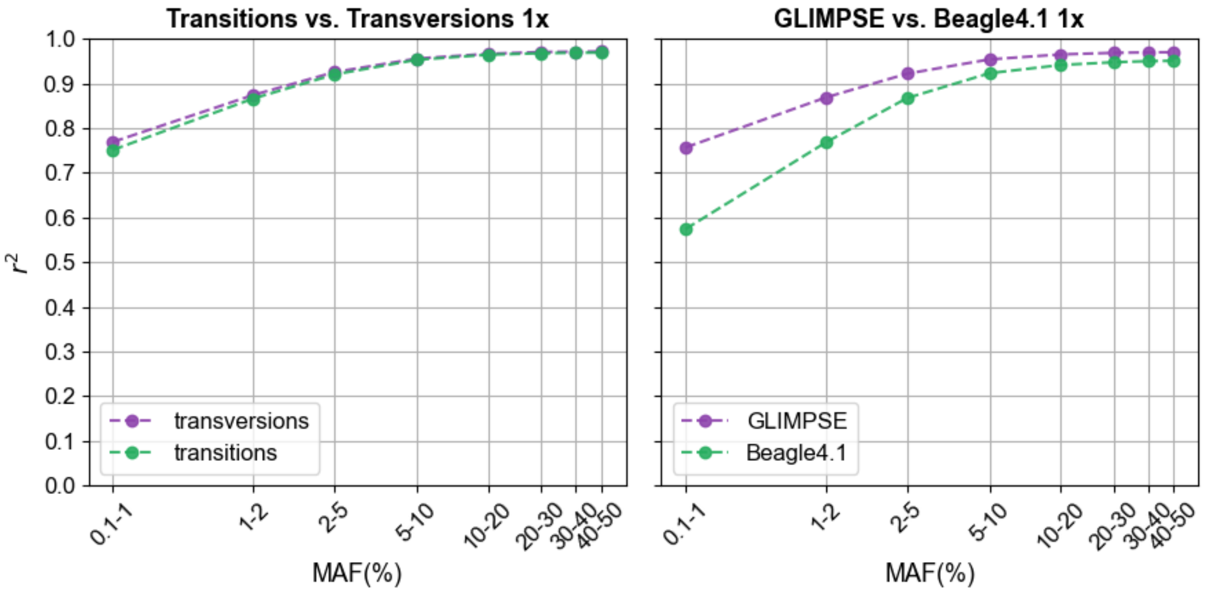

Figure S3: Imputation accuracy, $r^2$, for the aggregate of the 42 ancient genomes, previously downsampled to 1x, as a function of 1000 Genomes Project minor allele frequency (MAF), 0.1-50%, regarding (left to right) i) transitions (green) compared to transversions (purple), and ii) comparison between imputation methods, Beagle4.1[24] (green) vs. GLIMPSE1.1.1[25] (purple).

### 5. PCA with focus on Europeans, several coverages

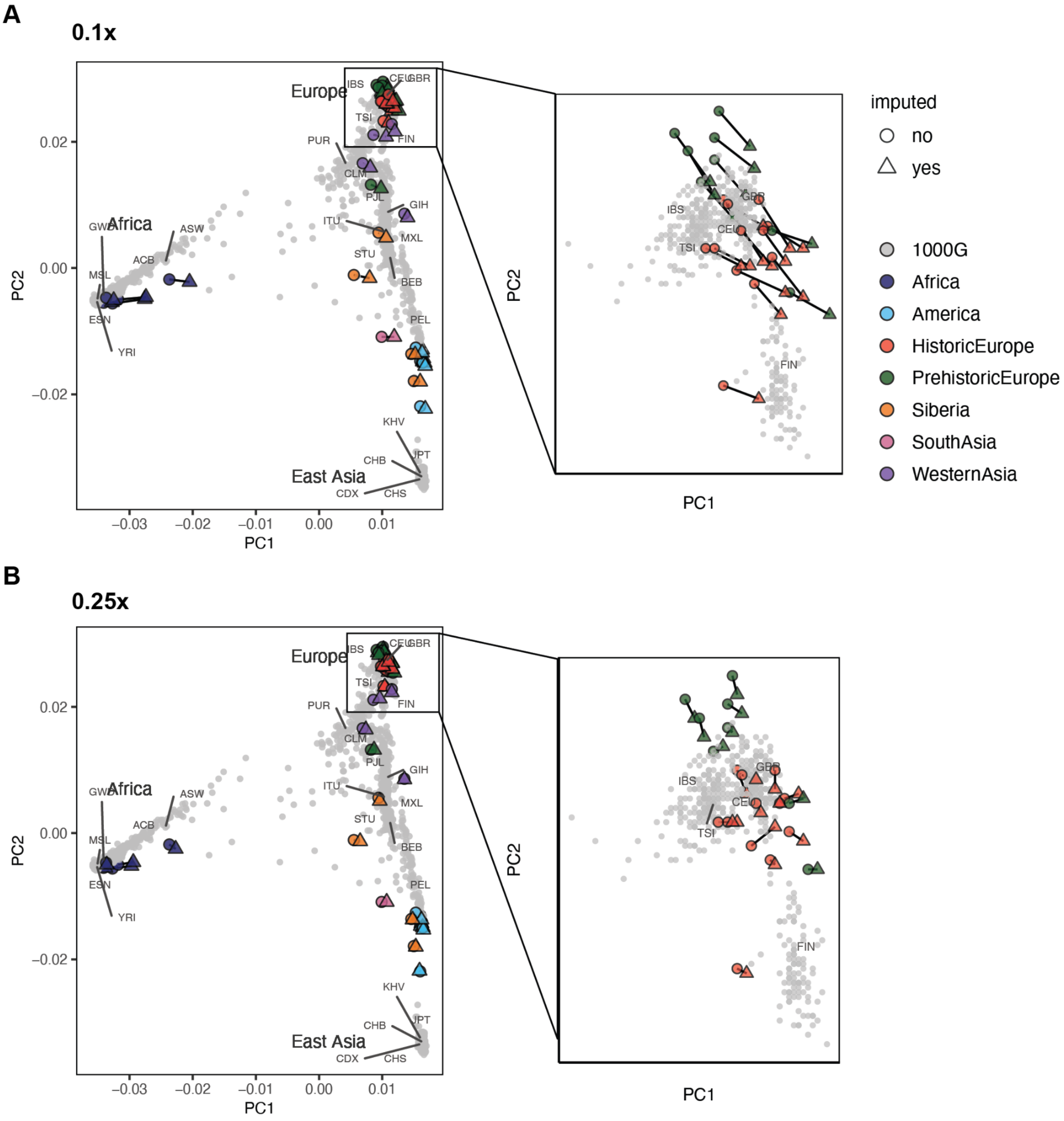

Figure S4: Principal component analysis (PCA) of imputed 0.1x (A) and 0.25x (B) and high-coverage ancient genetic data, and present-day data in 1000 Genomes reference panel (gray). Plots show individual coordinates along the two first principal components, zooming-in on the individuals of European ancestry. Imputed data points are represented by triangles and high-coverage ancient data by full circles. Corresponding imputed and high-coverage data are connected by a line.

154 ## 6. Genetic clustering analyses: from K=2 to K=5 clustering populations

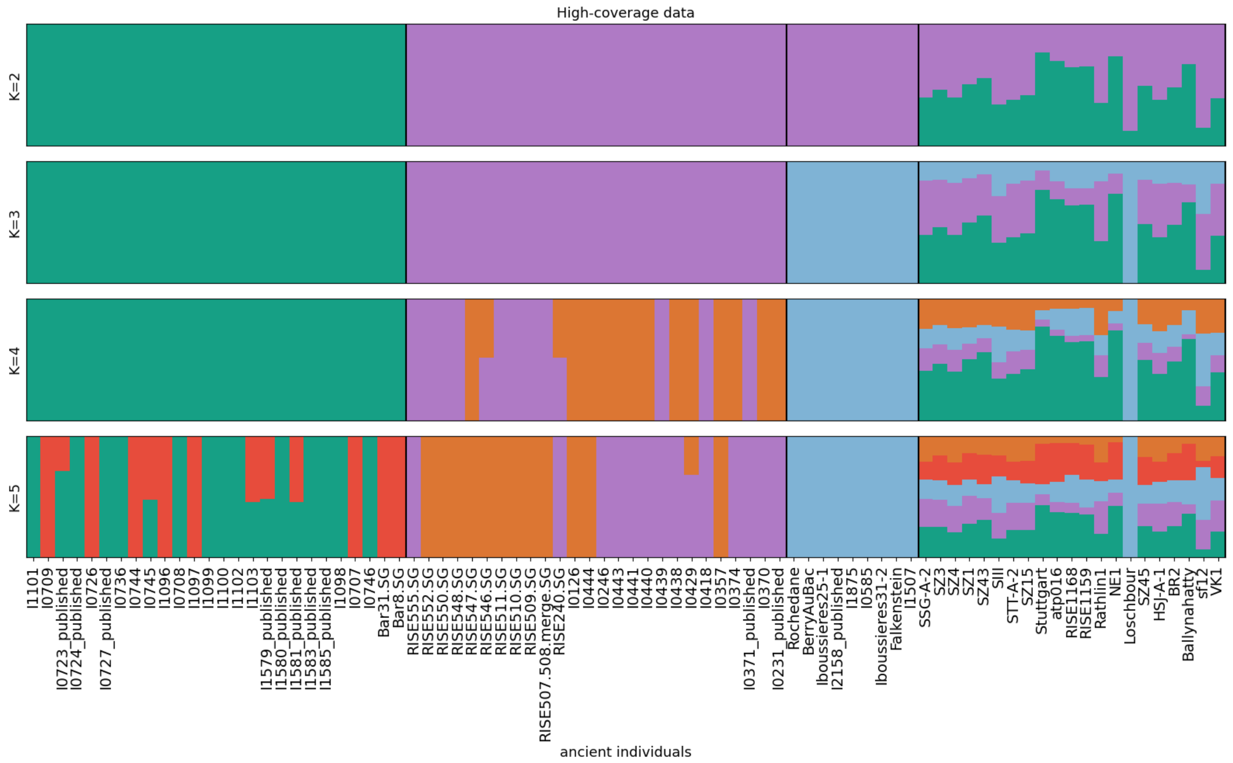

155

156 Figure S5: Genetic clustering results obtained from running unsupervised ADMIXTURE[26] with the

157 genetic data of a subset of individuals in 1240K dataset[27] and high-coverage data (rightmost box).

158

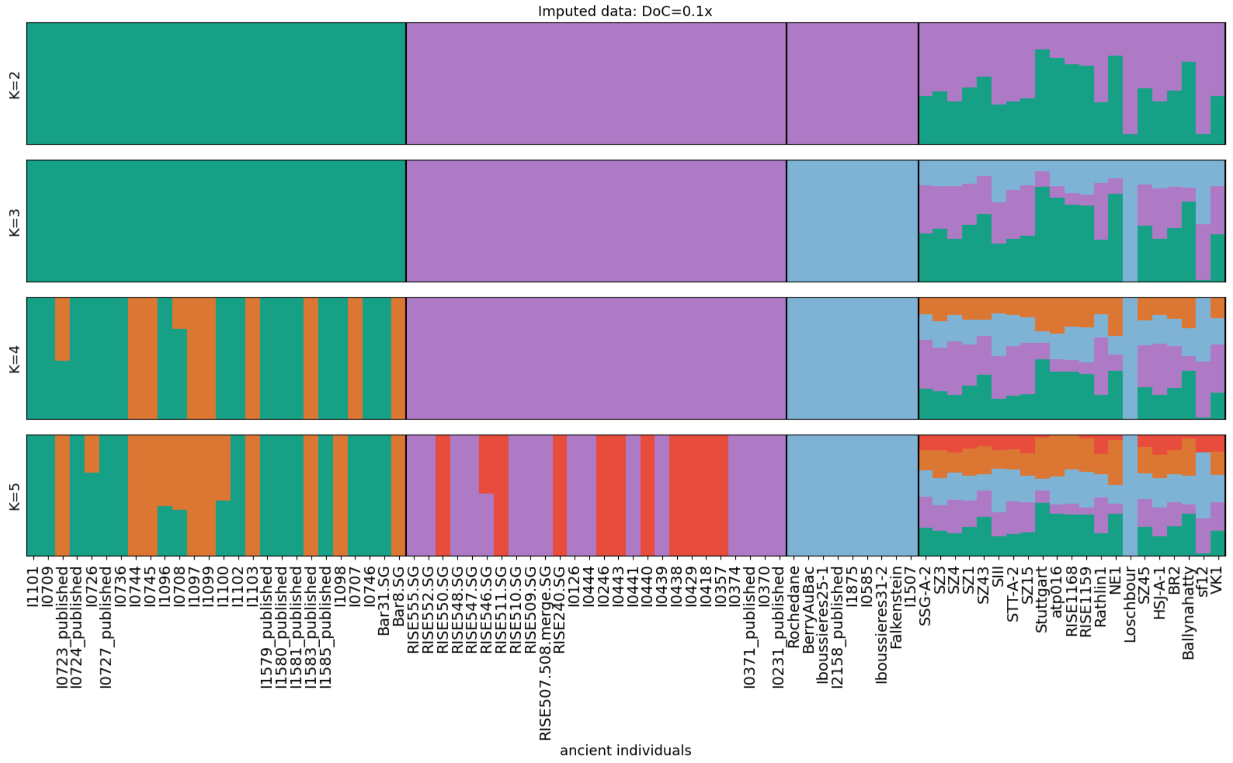

159

Figure S6: Genetic clustering results obtained from running unsupervised ADMIXTURE with the genetic data of a subset of individuals in 1240K dataset and imputed 0.1x data (rightmost box).

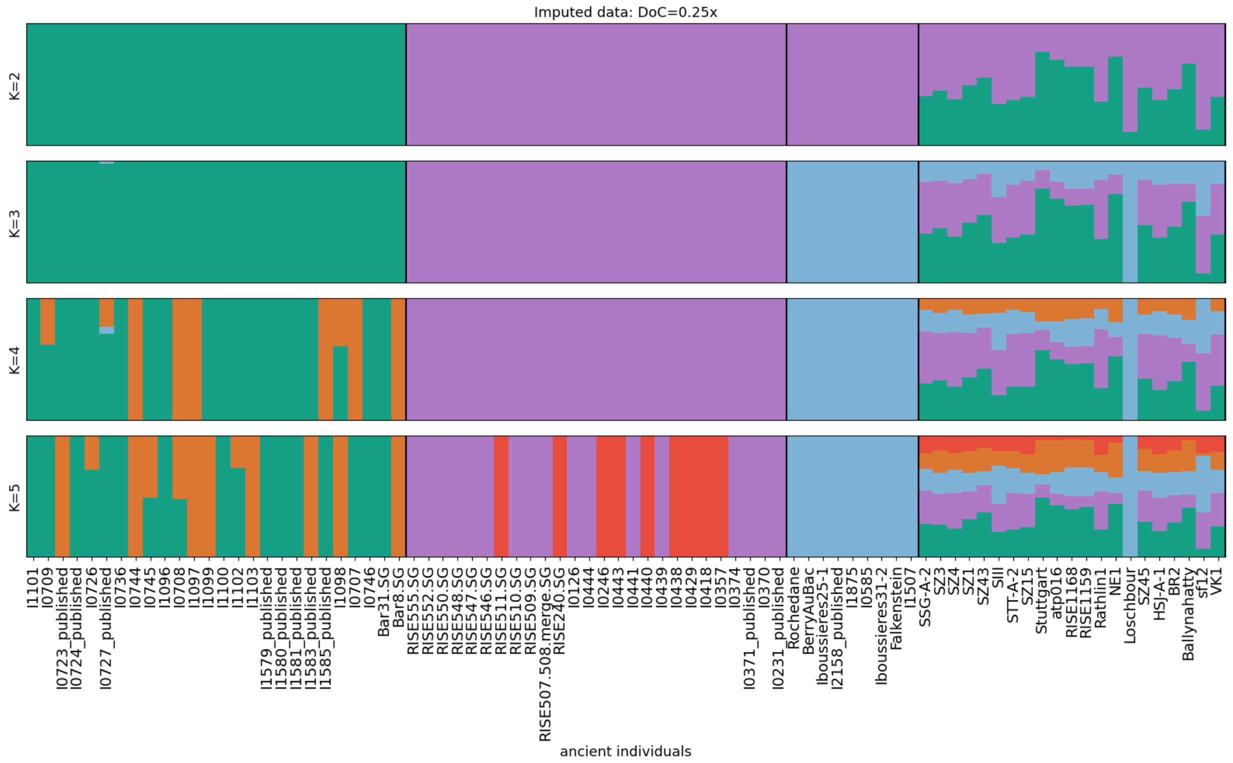

Figure S7: Genetic clustering results obtained from running unsupervised ADMIXTURE with the genetic data of a subset of individuals in 1240K dataset and imputed 0.25x data (rightmost box).

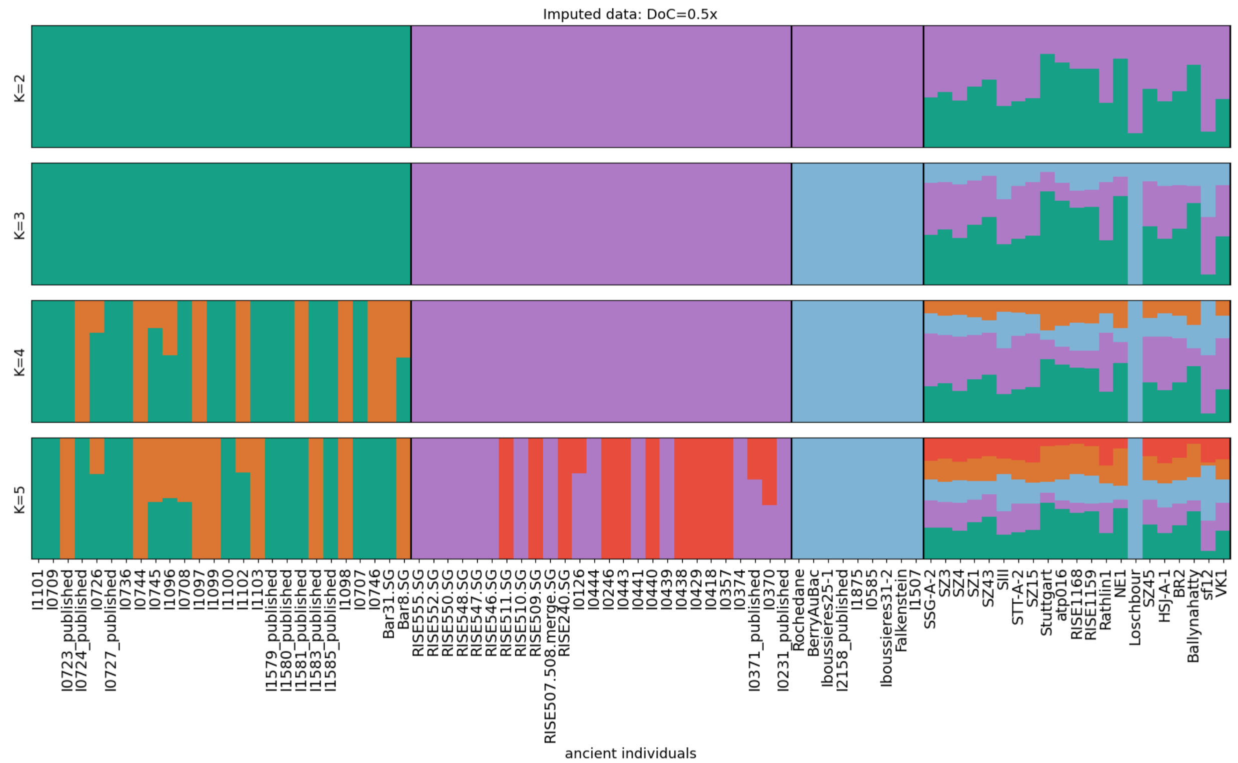

Figure S8: Genetic clustering results obtained from running unsupervised ADMIXTURE with the genetic data of a subset of individuals in 1240K dataset and imputed 0.5x data (rightmost box).

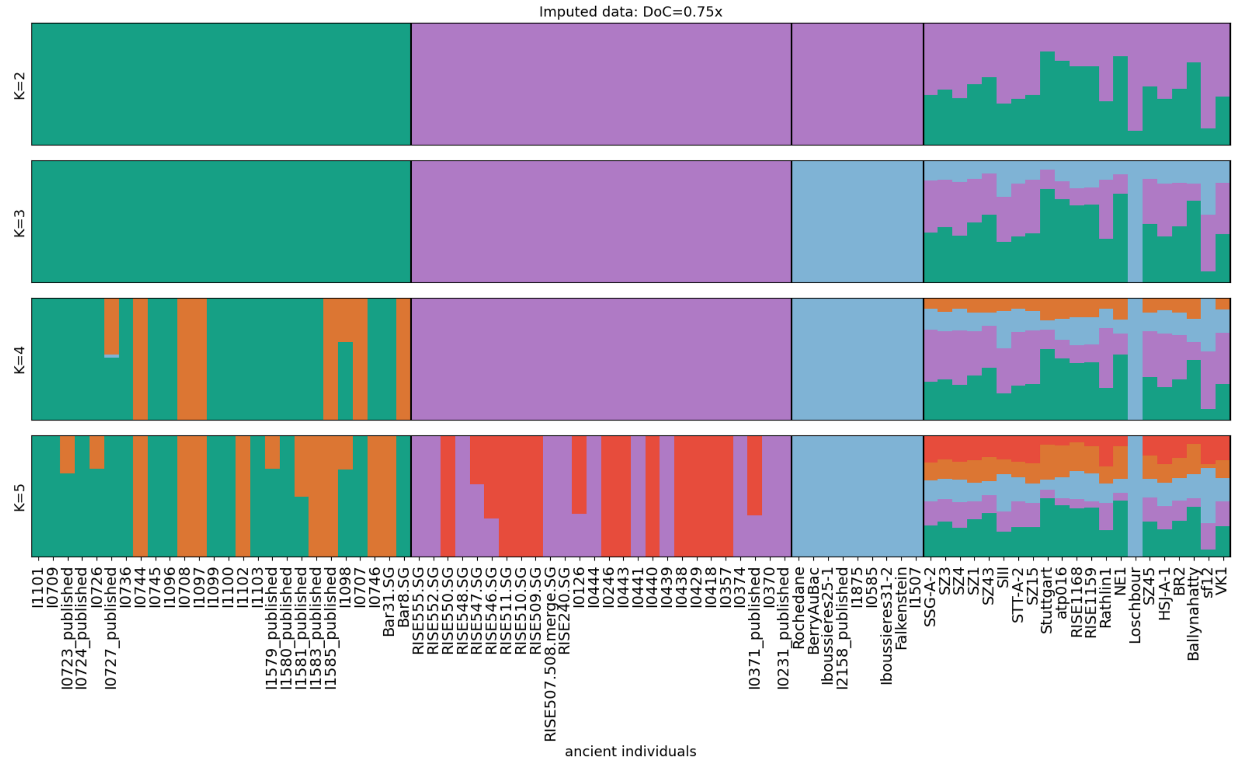

Figure S9: Genetic clustering results obtained from running unsupervised ADMIXTURE with the genetic data of a subset of individuals in 1240K dataset and imputed 0.75x data (rightmost box).

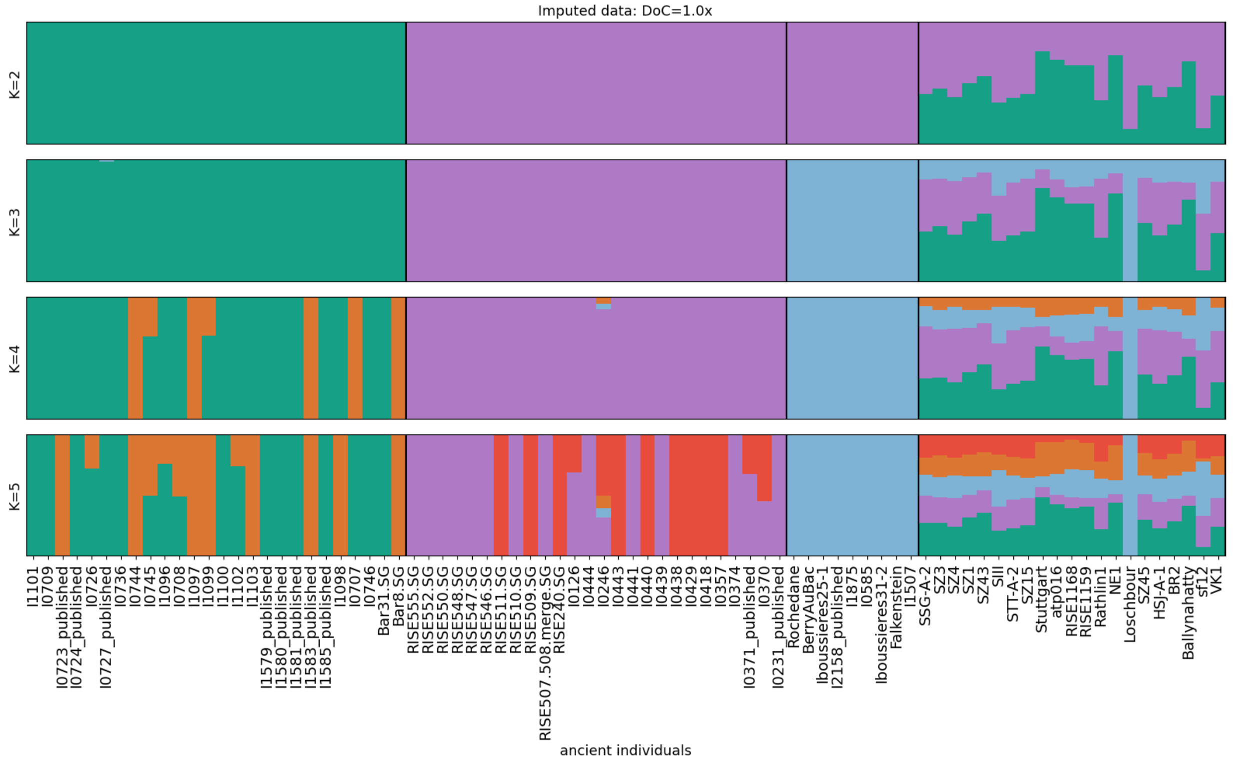

Figure S10: Genetic clustering results obtained from running unsupervised ADMIXTURE with the genetic data of a subset of individuals in 1240K dataset and imputed 1.0x data (rightmost box).

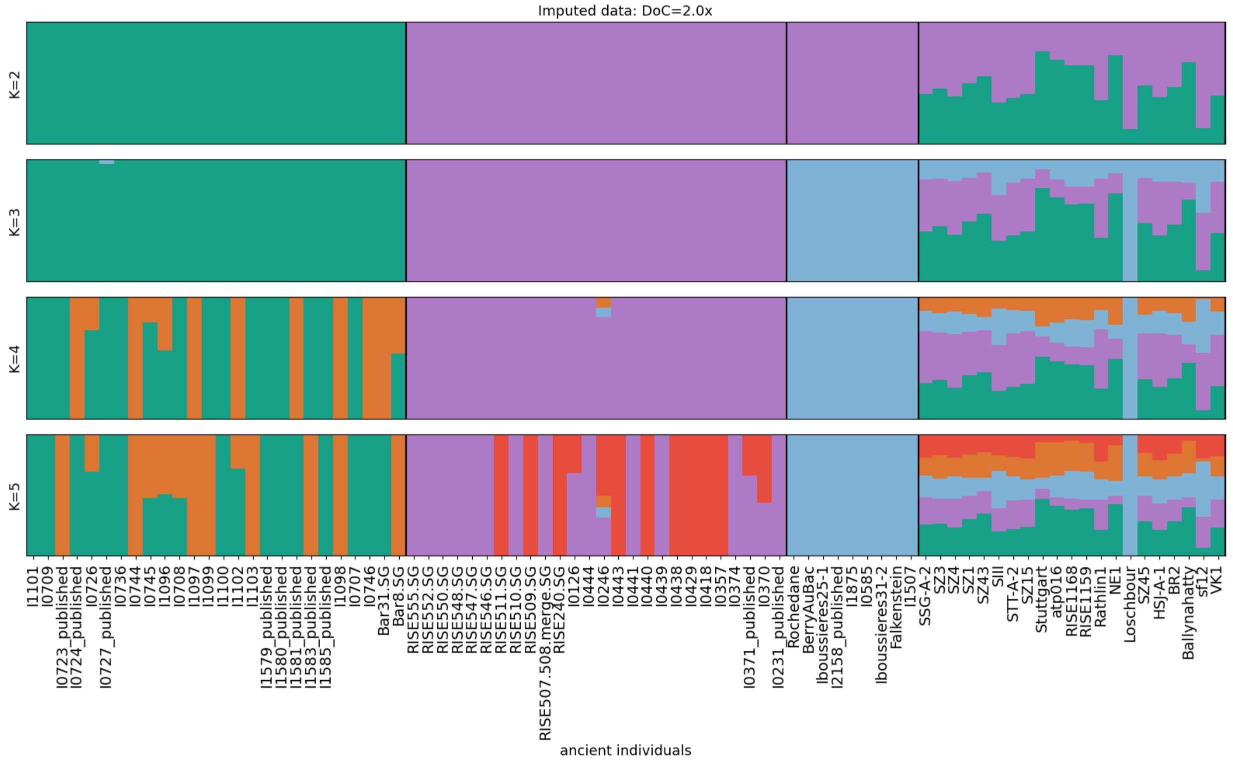

Figure S11: Genetic clustering results obtained from running unsupervised ADMIXTURE with the genetic data of a subset of individuals in 1240K dataset and imputed 2.0x data (rightmost box).

Table S2: Subset of the 1240K dataset[27] used as reference in the genetic clustering analyses. The individuals are grouped by population: Neolithic Anatolia (“Anatolia_N”), western hunter-gatherer (“WHG”), Early to Middle Bronze Age Steppe (“Steppe_EMBA”).

| Population | Individual |
|---|---|
| Anatolia_N | Bar31.SG |
| Anatolia_N | Bar8.SG |
| Anatolia_N | I0707 |
| Anatolia_N | I0708 |
| Anatolia_N | I0709 |
| Anatolia_N | I0723_published |
| Anatolia_N | I0724_published |
| Anatolia_N | I0726 |
| Anatolia_N | I0727_published |
| Anatolia_N | I0736 |
| Anatolia_N | I0744 |
| Anatolia_N | I0745 |
| Anatolia_N | I0746 |
| Anatolia_N | I1096 |
| Anatolia_N | I1097 |
| Anatolia_N | I1098 |
| Anatolia_N | I1099 |
| Anatolia_N | I1100 |
| Anatolia_N | I1101 |
| Anatolia_N | I1102 |
| Anatolia_N | I1103 |
| Anatolia_N | I1579_published |
| Anatolia_N | I1580_published |
| Anatolia_N | I1581_published |
| Anatolia_N | I1583_published |
| Anatolia_N | I1585_published |
| WHG | BerryAuBac |
| WHG | Falkenstein |
| WHG | I0585 |
| WHG | I1507 |
| WHG | I1875 |
| WHG | I2158_published |
| WHG | Iboussieres25-1 |
| WHG | Iboussieres31-2 |
| WHG | Rochedane |
| Steppe_EMBA | I0126 |
| Steppe_EMBA | I0231_published |
| Steppe_EMBA | I0246 |
| Steppe_EMBA | I0357 |
| Steppe_EMBA | I0370 |
| Steppe_EMBA | I0371_published |
| Steppe_EMBA | I0374 |
| Steppe_EMBA | I0418 |
| Steppe_EMBA | I0429 |
| Steppe_EMBA | I0438 |
| Steppe_EMBA | I0439 |
| Steppe_EMBA | I0440 |
| Steppe_EMBA | I0441 |
| Steppe_EMBA | I0443 |
| Steppe_EMBA | I0444 |
| Steppe_EMBA | RISE240.SG |
| Steppe_EMBA | RISE507.508.merge.SG |
| Steppe_EMBA | RISE509.SG |
| Steppe_EMBA | RISE510.SG |
| Steppe_EMBA | RISE511.SG |
| Steppe_EMBA | RISE546.SG |
| Steppe_EMBA | RISE547.SG |
| Steppe_EMBA | RISE548.SG |
| Steppe_EMBA | RISE550.SG |
| Steppe_EMBA | RISE552.SG |
| Steppe_EMBA | RISE555.SG |

### 7. ROH estimates for Sumidouro5

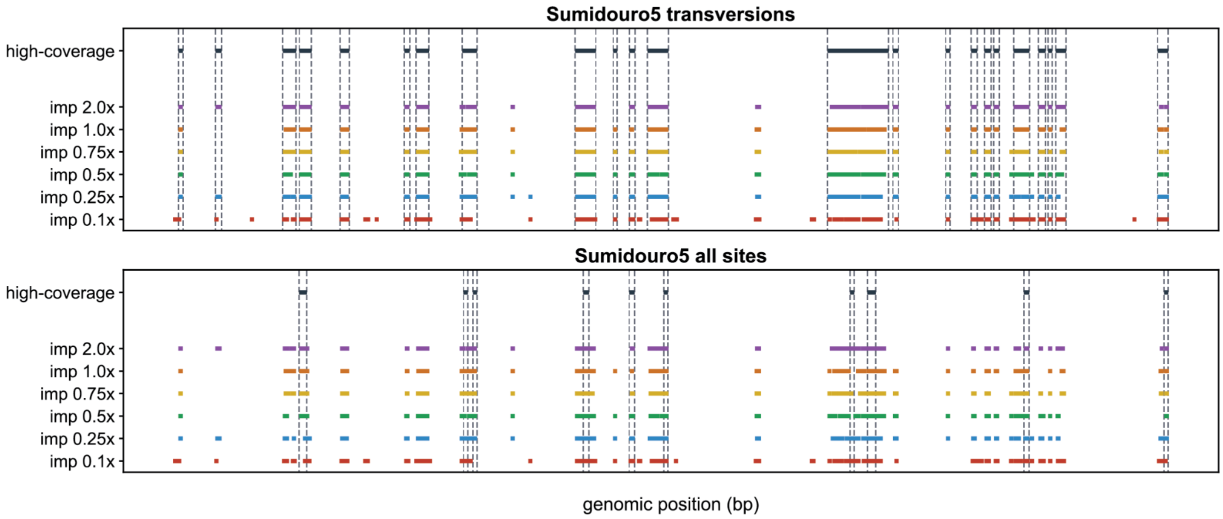

Figure S12: ROH segments identified in chromosome 10 for high-coverage and imputed data (DoC between 0.1x and 2.0x) for Sumidouro5. Top: ROH obtained using all sites. Bottom: ROH estimated using transversion sites only.
